# An update on the mosquito fauna of Aguascalientes, Mexico: morphological and molecular identifications, host-feeding patterns, and the removal of *Aedes idahoensis* (Theobald) from Mexico

**DOI:** 10.64898/2026.10.13.757877

**Authors:** Aldo I. Ortega-Morales, Nadia A. Fernández-Santos, Isis J. Morales-Avitia, Guadalupe Flores-Salas, Lihua Wei, Miguel Medrano-Santillana, Sujata Balasubramanian, Sarah A. Hamer, Gabriel L. Hamer, Herón Huerta, Jessica M. Flores-Salas, Jorge J. Rodríguez-Rojas, Mario A. Rodríguez-Perez

**Affiliations:** Departamento de Parasitología, Universidad Autónoma Agraria Antonio Narro, unidad Laguna, Torreón, Coahuila, Mexico; Instituto Politécnico Nacional, Centro de Biotecnología Genómica, Reynosa, Tamaulipas, Mexico; Department of Entomology, Texas A&M University, College Station, Texas, United States of America; Programa de Posgrado en Producción Agropecuaria, Universidad Autónoma Agraria Antonio Narro, unidad Laguna, Torreón, Coahuila, Mexico; Departamento de Biología, Universidad Autónoma Agraria Antonio Narro, unidad Laguna, Torreón, Coahuila, Mexico; Department of Veterinary Integrative Biosciences, Texas A&M University, College Station, Texas, United States of America; Instituto de Diagnóstico y Referencia Epidemiológicos, Secretaria de Salud, Mexico City, Mexico; Departamento de Ciencias Medico Veterinarias, Universidad Autónoma Agraria Antonio Narro, unidad Laguna, Torreón, Coahuila, Mexico; Facultad de Agronomía, Campus de Ciencias Agropecurarias, Laboratorio de Entomología, Universidad Autónoma de Nuevo León, Escobedo, Nuevo León, Mexico

## Abstract

To better understand taxonomic identity and feeding patterns of mosquitoes in all physiographic regions and sub-regions of Aguascalientes, Mexico, we conducted surveillance followed by morphologic analysis and DNA metabarcoding. We targeted the Sierra Madre Occidental, Central Plateau, and Neo-Volcanic Axis. We sampled natural larval habitat for immature stages and adults were collected using Shannon and CDC light traps, mechanical aspirators, and human landing catch method. In addition, curated specimens from Aguascalientes deposited in a local institution were re-examined. Overall, 5,227 mosquito specimens consisting of 357 larvae, 199 larval exuviae, 373 pupal exuviae, 33 pupae, 3,000 females, 1,264 males, and 1 male genitalia were examined using taxonomic keys. Most immature and adult species (*n* = 20/34 species representing 58.8% coverage) were barcoded for molecular species designation, and a subsample (*n* = 64) of engorged female mosquitoes were analyzed for blood feeding patterns. Two subfamilies, four tribes, seven genera, 14 subgenera, and 34 species were identified, which one tribe, three genera, nine subgenera, and 22 species are new records. Phylogenetic analysis of specimens from Aguascalientes identified 20 species. Morphological observation of mosquitoes deposited in entomological collections resulted in the removal of *Aedes idahoensis* from Mexico, leaving the country with a total of 255 species documented. Bloodmeal metabarcoding was successful in 23 mosquitoes of five species, revealing evidence of feeding on more than a single vertebrate species in 17 individuals. Birds were the most common host with the cattle egret (*Ardea ibis*) found in 18 out of 23 mosquitoes. This update on the taxonomic identity and mosquito richness and feeding patterns from Aguascalientes affords a better understanding of the ecology of medically important mosquito species.

**Author summary:** While mosquito-borne diseases are a major public health problem in Mexico, many Mexican states have received minimal attention to documenting the mosquito communities. We conducted a comprehensive survey of mosquito communities in three regions of the state of Aguascalientes, located in central Mexico. We sampled natural habitats for immature mosquitoes and adult mosquitoes were collected by traps and with aspirators. In addition, curated museum specimens from Aguascalientes were re-examined. Overall, 5,227 mosquito specimens consisting of 3,000 adult females, 1,264 adult males, 357 larvae, 199 larval exuviae, 373 pupal exuviae, 33 pupae, and 1 male genitalia were examined using taxonomic keys. Two subfamilies, four tribes, seven genera, 14 subgenera, and 34 species were identified; one tribe, three genera, nine subgenera, and 22 species are new state records. A subset of 20 taxa were barcoded for species identification. Morphological re-examination of museum collections resulted in the removal of *Aedes idahoensis* from Mexico, leaving the country with a total of 255 species documented. A subsample of 23 engorged female mosquitoes of five species were analyzed for blood feeding patterns using bloodmeal metabarcoding revealing evidence of feeding on more than a single vertebrate species in 17 individuals. Birds were the most common host with the cattle egret (*Ardea ibis*) found in 18 out of 23 mosquitoes. This study fills knowledge gaps on the mosquito community and host contact in Aguscalientes, a neglected state of Mexico.

## Introduction

In Mexico, mosquito-borne diseases are a significant public health problem, with continued transmission of the pathogens that cause dengue fever, chikungunya and Zika, among others. Other mosquito-borne diseases such as yellow fever, Mayaro, and various encephalitis are underestimated. More than 50 million people are at risk of contracting one of these diseases, where active transmission cycles are maintained in most of the national territory [1].

Mexico is divided in 32 states, with 12 receiving mosquito surveillance focused on taxonomy, ecology, and spatial occurrence. Lower taxa richness of 27 and 30 mosquito species are reported from Tlaxcala and Mexico City, respectively [2–6]; higher richness, ranging from 50–73 mosquito species per state have been reported from Querétaro [7–8], Mexico State [9], Sonora [10], Hidalgo [11–13], Morelos [14], and Nuevo León [15–20]. The highest richness of mosquito species includes 78, 84, 107, and 142 species, reported from Quintana Roo [21–30], Tamaulipas [31–32], Tabasco [33–35], and Veracruz [36–39], respectively.

In the state of Aguascalientes, dengue fever is the most important mosquito-borne disease, where it is endemic [40]. Other diseases such as chikungunya and Zika have been also reported in Aguascalientes state recently [41–42]. However, mosquito communities of Aguascalientes are poorly characterized, as only 12 mosquito species have been reported with most records based on mosquito surveillance studies from urban and sub-urban regions. Here, a more comprehensive mosquito surveillance study was conducted, which includes the three physiographical regions of Aguascalientes, namely, the “Sierra Madre Occidental”, “Central Plateau”, and “Neo-Volcanic Axis” [43]. Mosquito surveillance included remote forest and mountain regions in dry and rainy seasons.

The identification of bloodmeal hosts of mosquitoes provides essential ecological insight into mosquito-host contact. Different mosquito taxa are known to have preferences for different vertebrate hosts, and these intrinsic factors directly influence the transmission potential of multiple disease-causing agents. Increasing appreciation for individual mosquitoes to take multiple bloodmeals from different host species within a single gonotrophic cycle [44–45] has implications for pathogen maintenance and transmission.

The objectives of this study were to characterize the mosquito taxonomic composition and ecology for populations in Aguascalientes. We combined morphological identification of adult and immature specimens with molecular barcoding to identify specimens. We used bloodmeal metabarcoding to document mosquito-host associations. This study fills important knowledge gaps about medically important mosquitoes of a neglected region of central Mexico.

## Materials and Methods

### Study Area

Aguascalientes is in the central part of Mexico, between 22° 27’ and 21° 38’north latitude and the meridians 101° 53’ and 102° 52’ west longitude. The state has an area of 5,680 km^2^, altitude of 1770-3050 m above sea level, and is a relatively small state in Mexico that is highly industrialized. It is bordered to the north and west by Zacatecas; and by Jalisco to the south and east. The state is further divided into three physiographic regions and three sub-regions (Fig 1). Namely, “Sierra Madre Occidental” (Zacatecas Mountains and Valleys), “Central Plateau” (Ojuelos-Aguascalientes Plateau), and “Neo-Volcanic Axis” (Highlands of Jalisco) [43]. A description of the regions and sub-regions of Aguascalientes, including the list of municipalities surveyed is given in Table 1.

**Fig 1.**
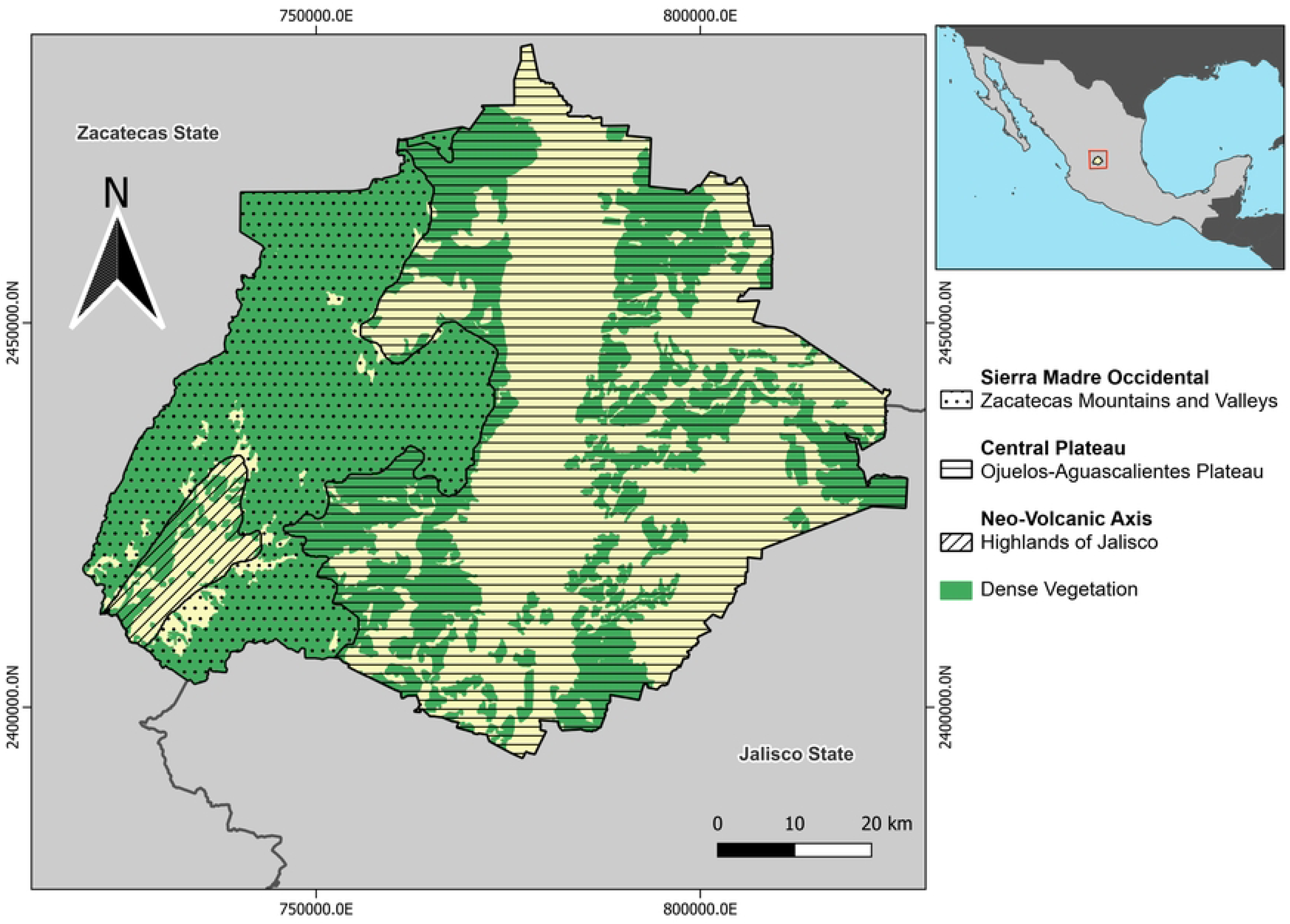
Physiographic map of the state of Aguascalientes, Mexico. Map showing the geographical boundaries and three physiographic regions with their respective sub-regions: Sierra Madre Occidental (Zacatecas Mountains and Valleys), Central Plateau (Ojuelos-Aguascalientes Plateau), and Neo-Volcanic Axis (Highlands of Jalisco).

**Table 1.** Description of the physiography of Aguascalientes, including the municipalities surveyed.

| Region (Sub-region) | Description of Sub-regions | Municipalities surveyed |
| --- | --- | --- |
| **Sierra Madre Occidental**<br>(Zacatecas Mountains and Valleys) | Mountainous regions with elevations greater than 2,500 masl. The climate is cold and cold most of the year. The vegetation consists mainly of oak and pine forests. | Calvillo, Jesús María, Rincón de Romos, San José de Gracia |
| **Central Plateau** (Ojuelos-Aguascalientes Plateau) | Extensive plains with elevations near to 2,000 masl, mostly covered with desert vegetation and grasslands. The climate is arid during the summer and cold during the winter. | Aguascalientes, Asientos, Cosío, El Llano, Pabellón de Arteaga, San Francisco de Romo, Tepezalá |
| **Neo-Volcanic Axis** (Highlands of Jalisco) | A large plateau with elevations near to 2,000 masl. It has a temperate dry and sub-humid climate. Includes extensive native grasslands and agricultural lands. | Aguascalientes |

### Mosquito Collection

Immature stages, except eggs, and adult mosquitoes were collected in 11 municipalities of the three physiographic regions of Aguascalientes (Table 1). The collections were conducted in both the dry (July) and rainy seasons (September–October) from 2023 through 2024. Immature stages were collected from all bodies of water serving as breeding sites. Larvae and pupae were placed alive in cups using water from the larval habitat and pH, salinity, and temperature recorded using a portable Hanna® (mod. HI98129) tester (Padua, Italy). Collected specimens were transported to a local lab. A portion of fourth-instar larvae from each collection was mounted on microscope slides using Euparal as mounting medium, whereas the rest of the live larvae were placed in individual emergence tubes to obtain adults with their associated larval and pupal exuviae. Male genitalia were dissected to assist identification when required. Host-seeking mosquitoes were collected using human landing catch method, Shannon traps containing protected human volunteers were deployed between 18:00 and 22:00 h and 10 Centers for Disease Control and Prevention (CDC) light traps deployed between 19:00 and 09:00 h. Resting mosquitoes were aspirated directly from caves and vegetation using mechanical aspirators. Adult mosquitoes were killed using triethylamine vapors and later mounted on insect pins. Mosquitoes mounted on insect pins were further identified using a stereomicroscope Zeiss® Discovery V8 (Oberkochen, Germany), while immature stages and exuviae identified using a microscope Zeiss® Primostar. Biological information such habitat, environmental parameters, and associated species for new records of Aguascalientes mosquitoes are shown in Table 2.

**Table 2.**
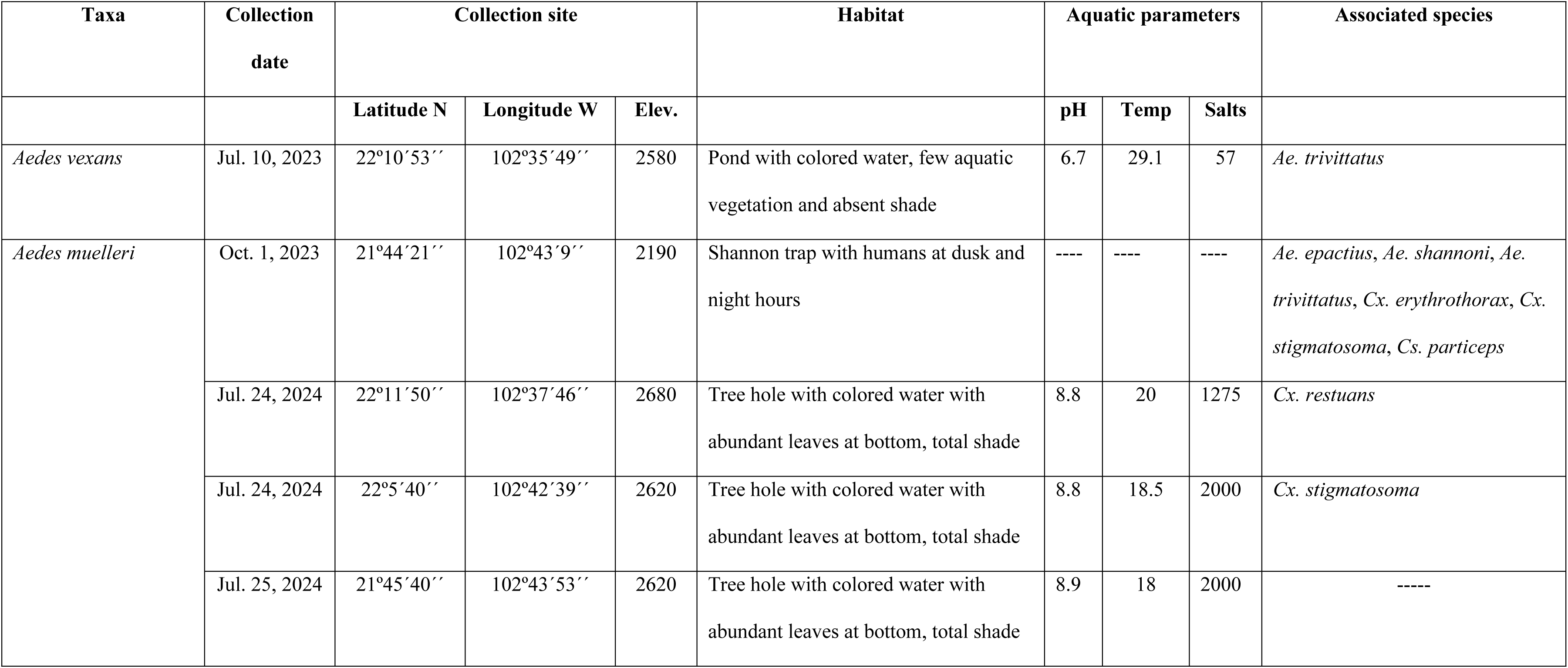
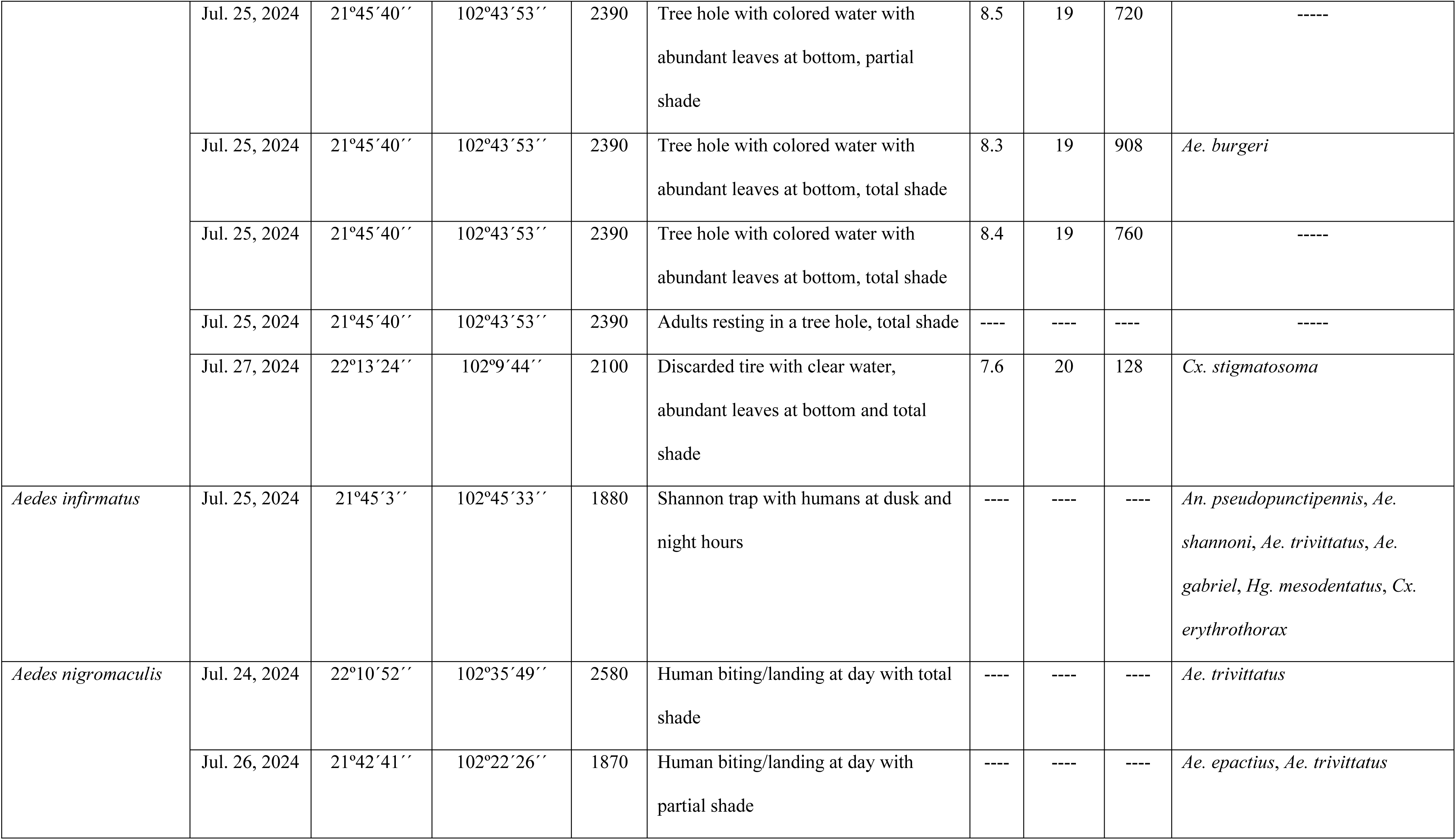
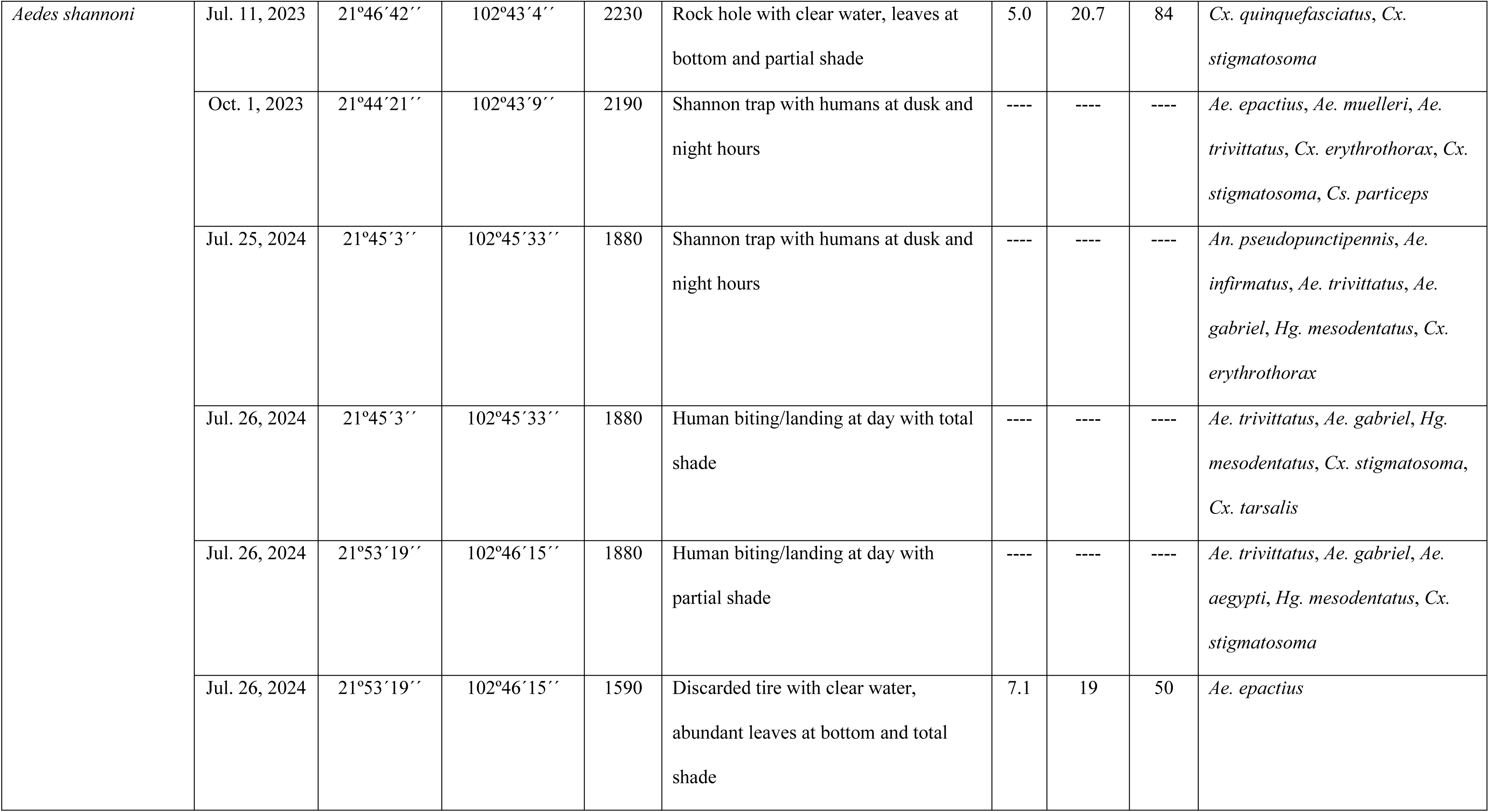
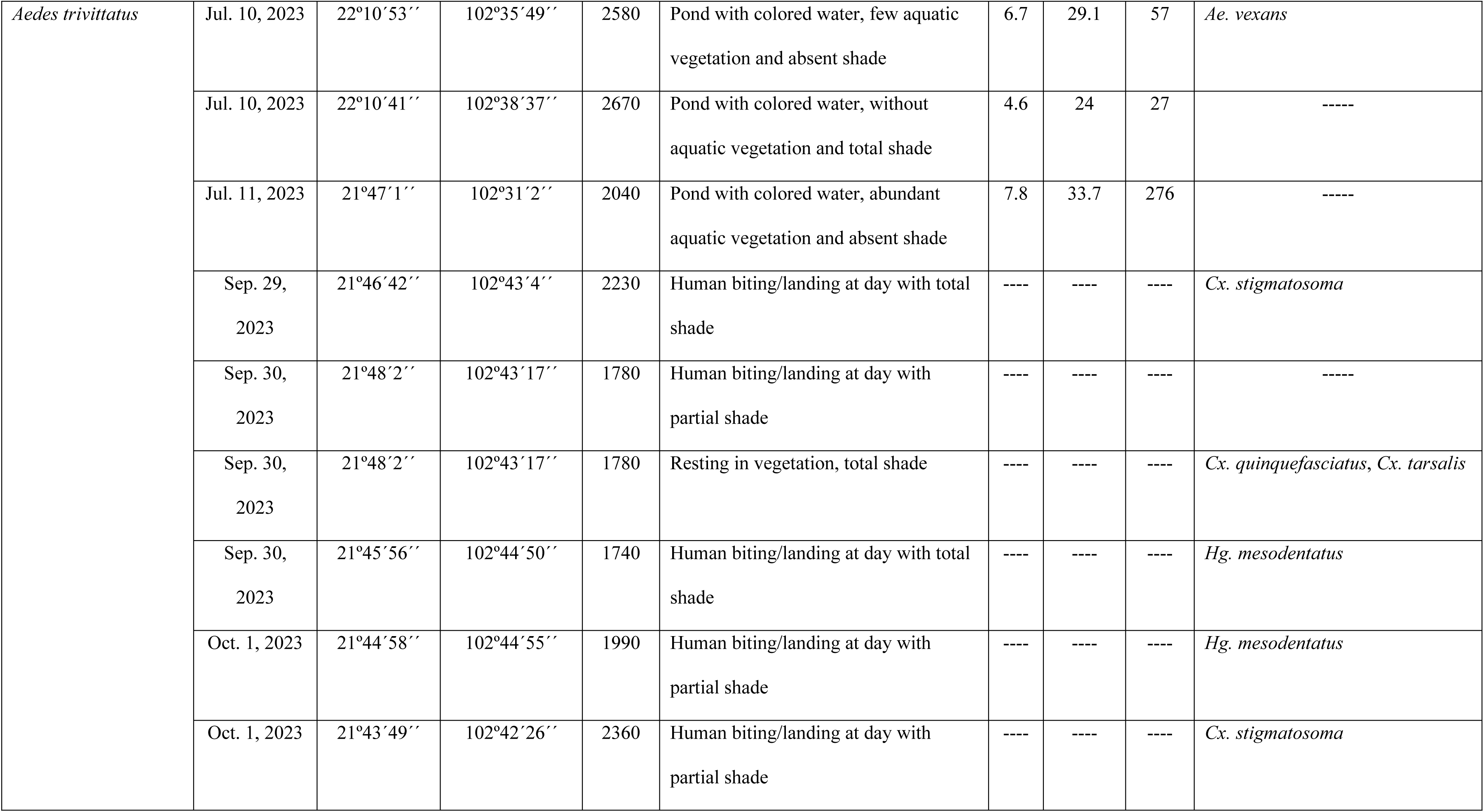
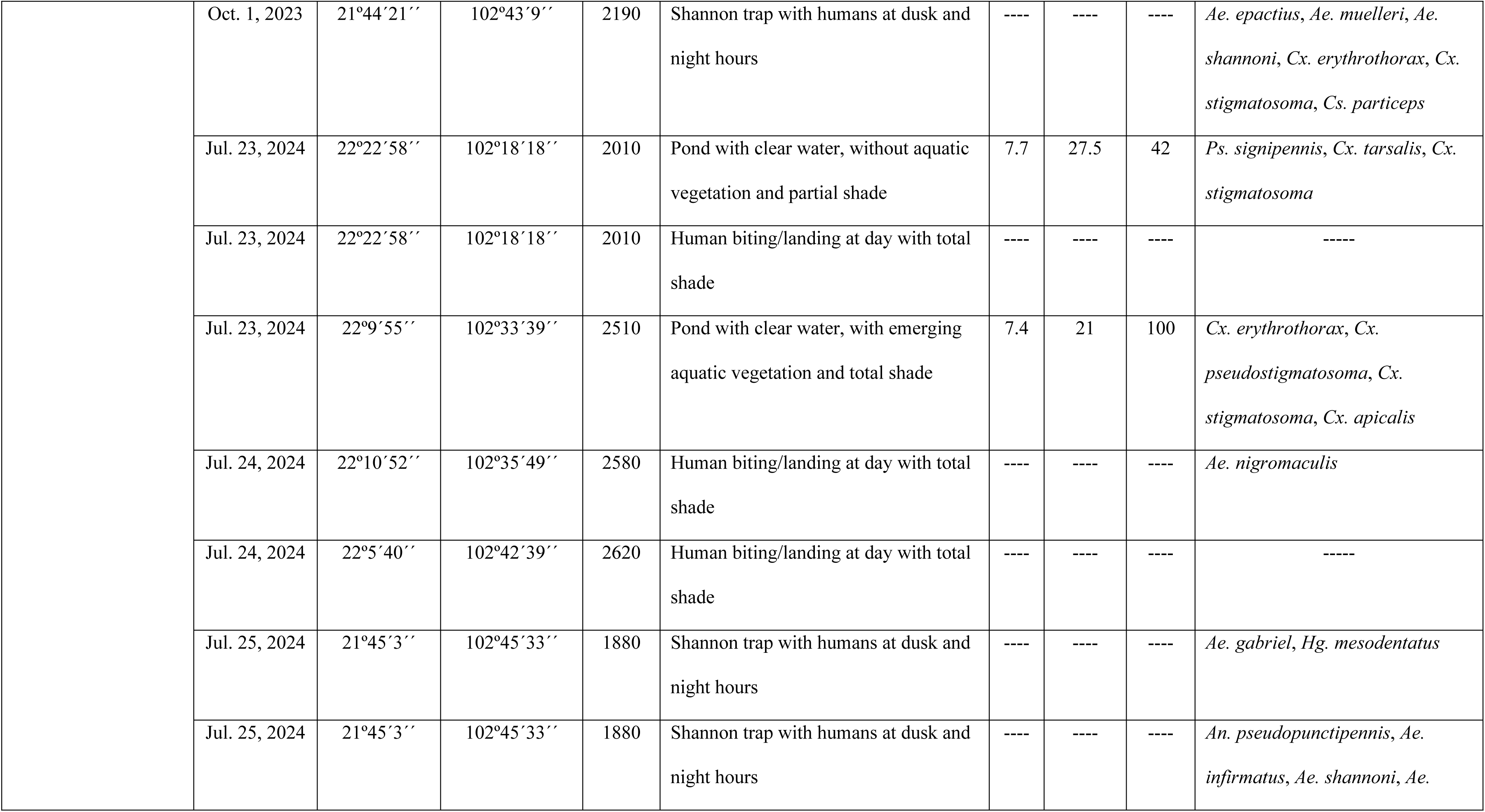
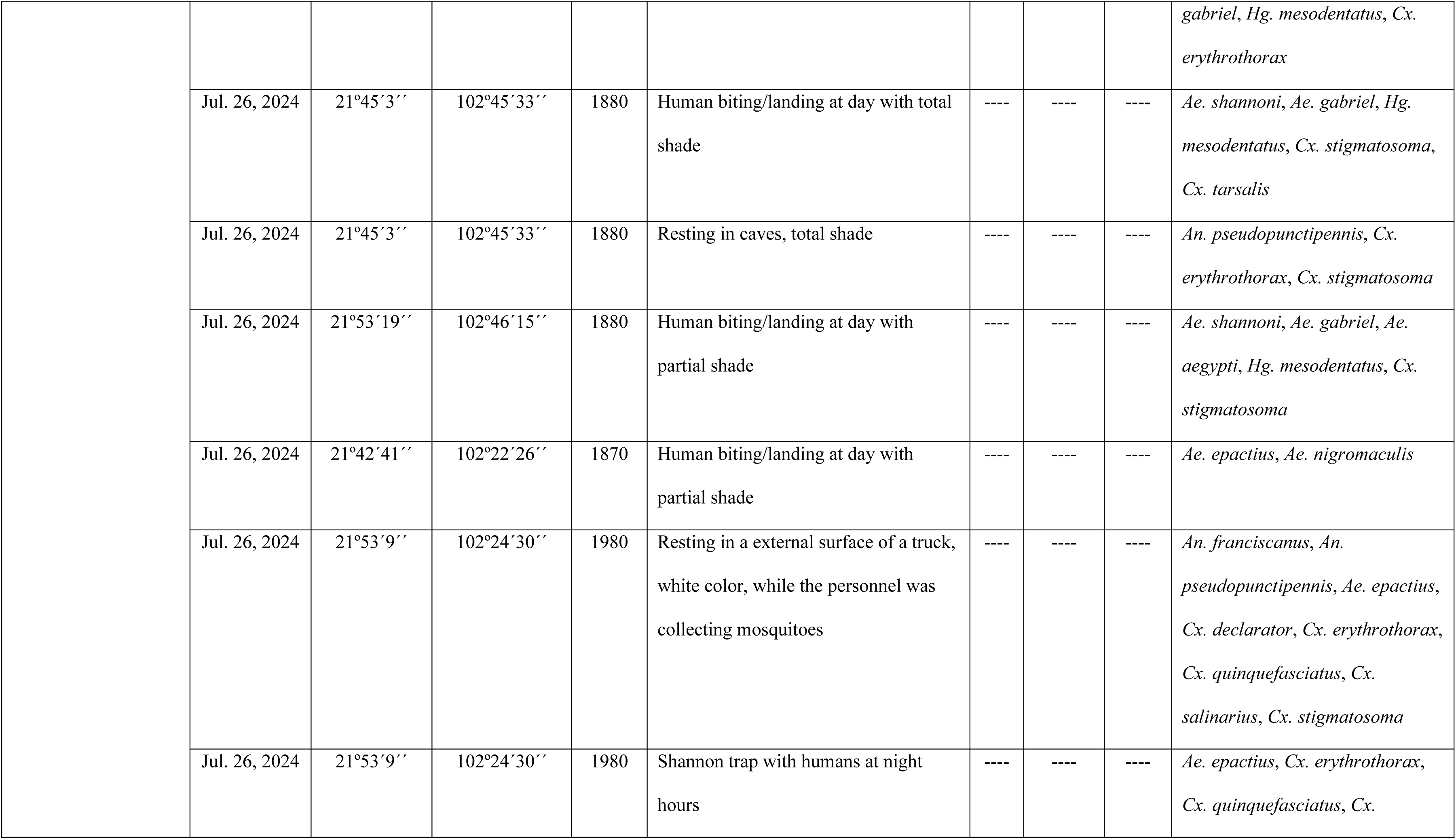
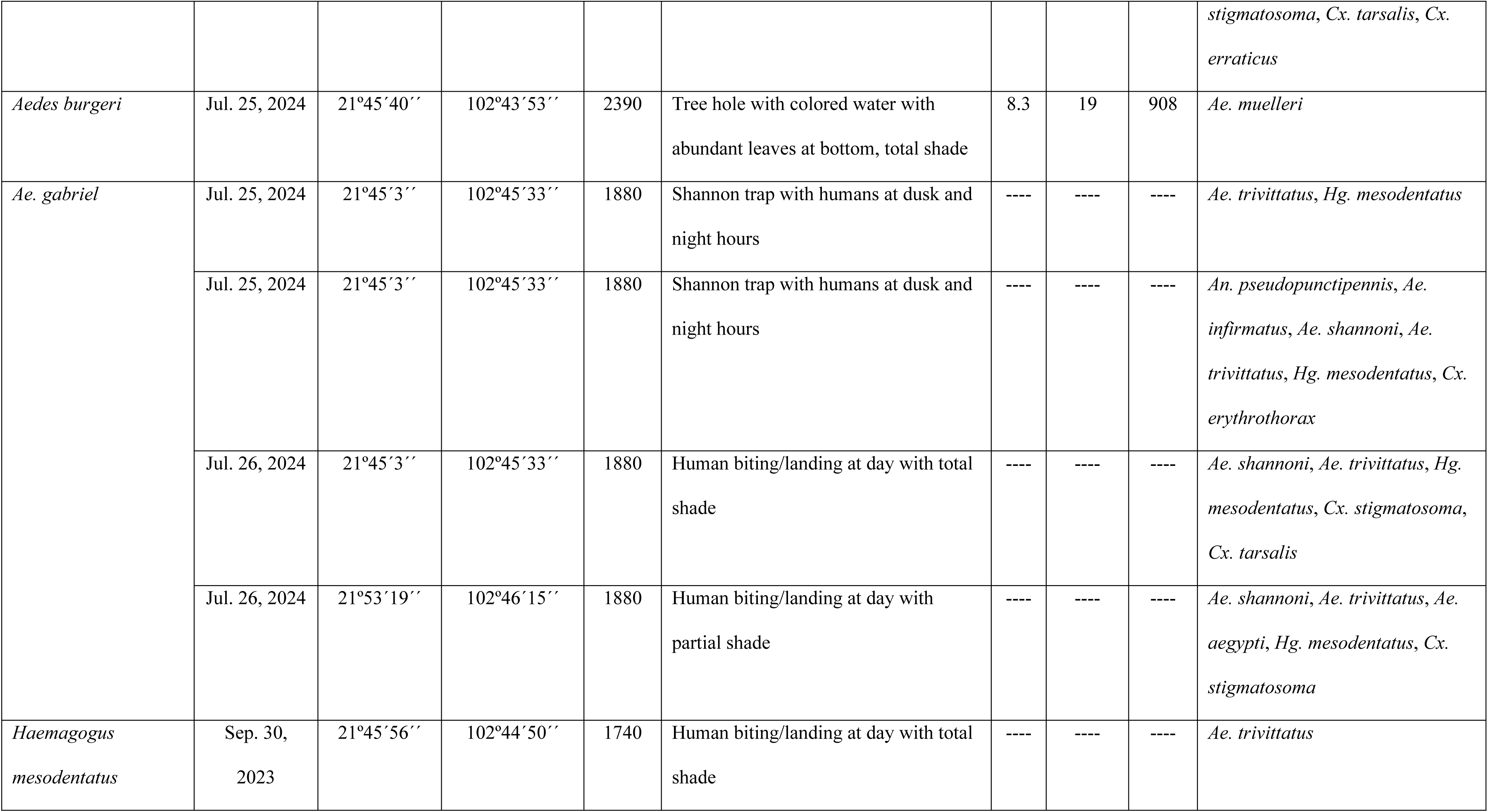
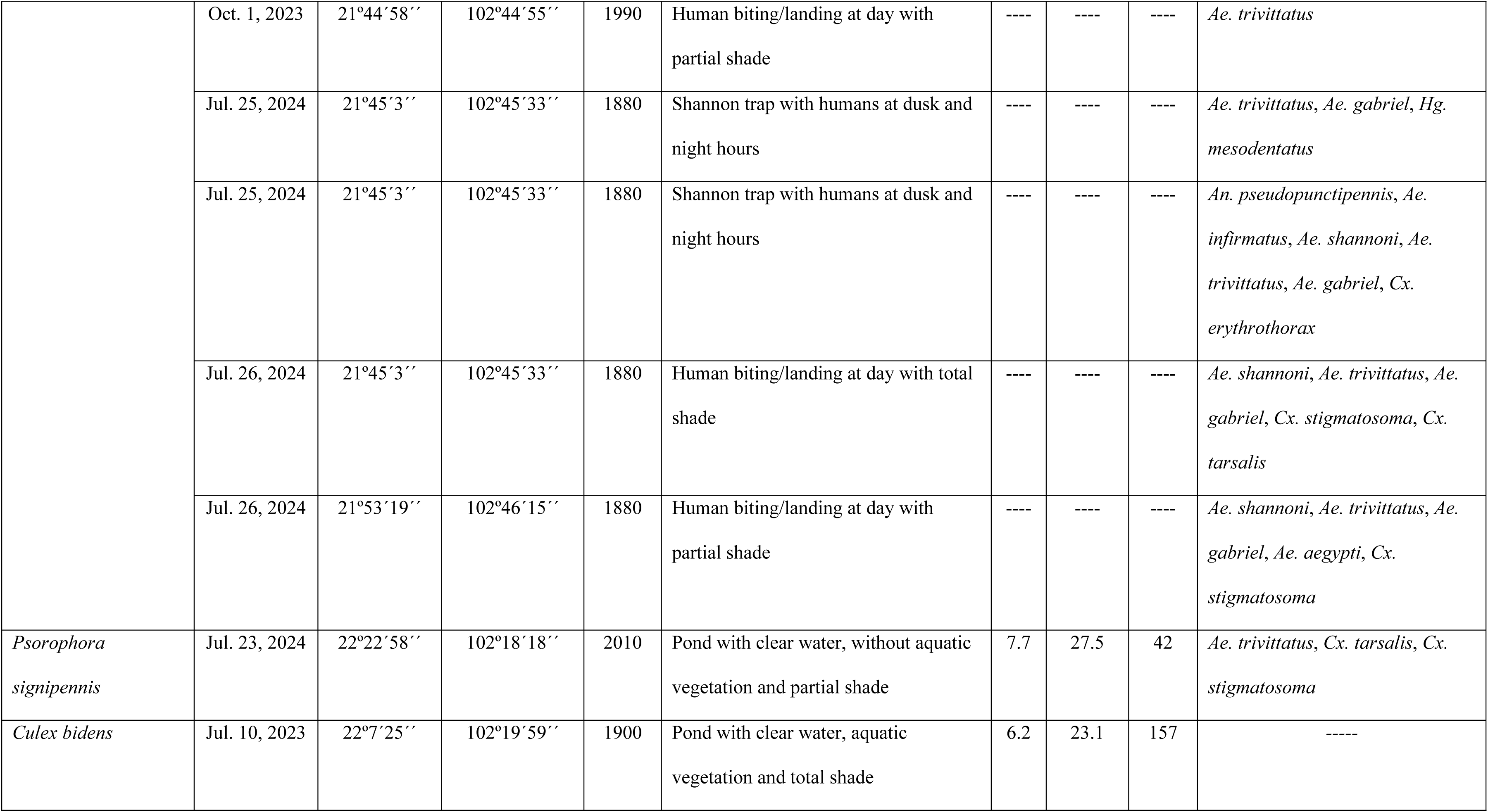
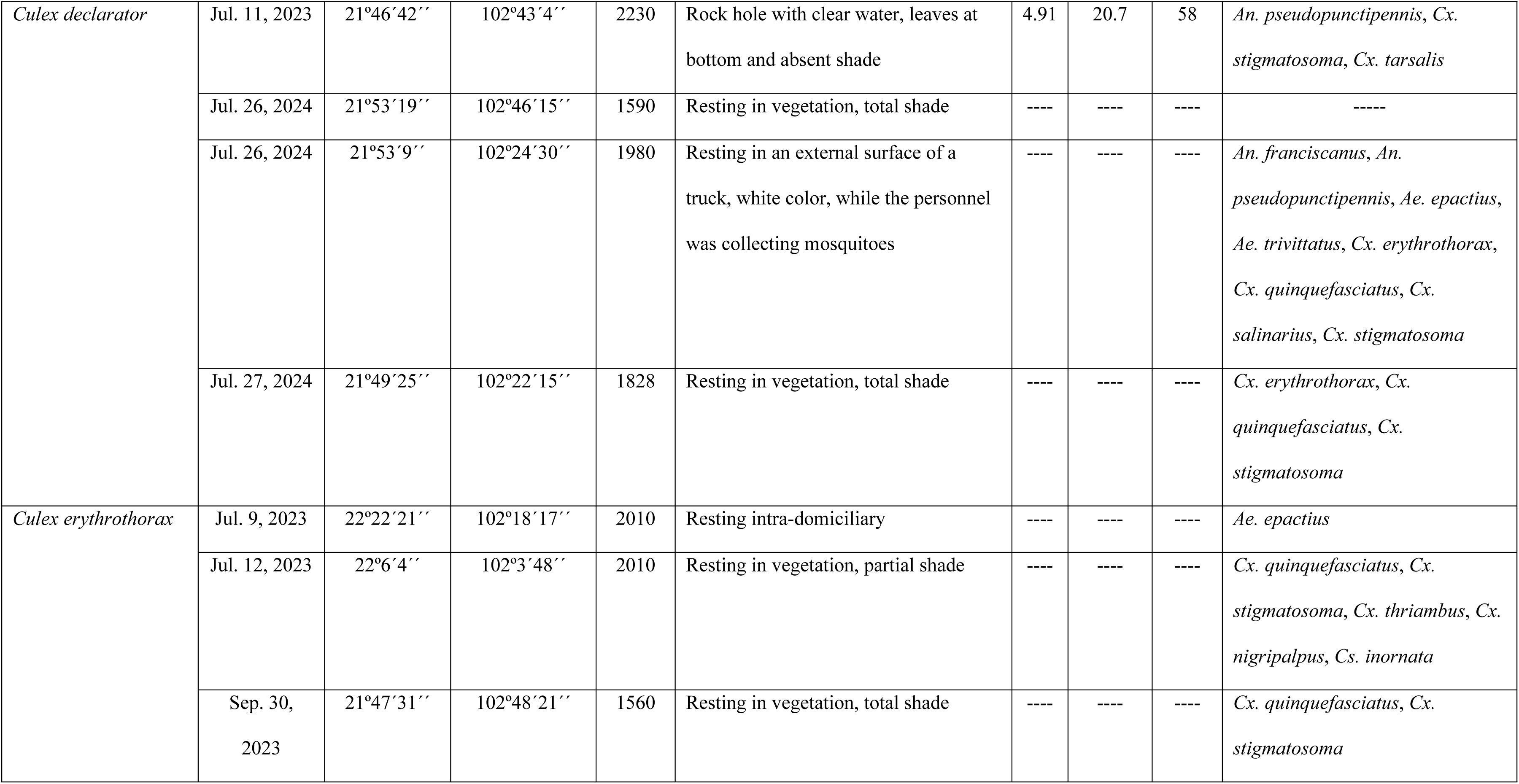
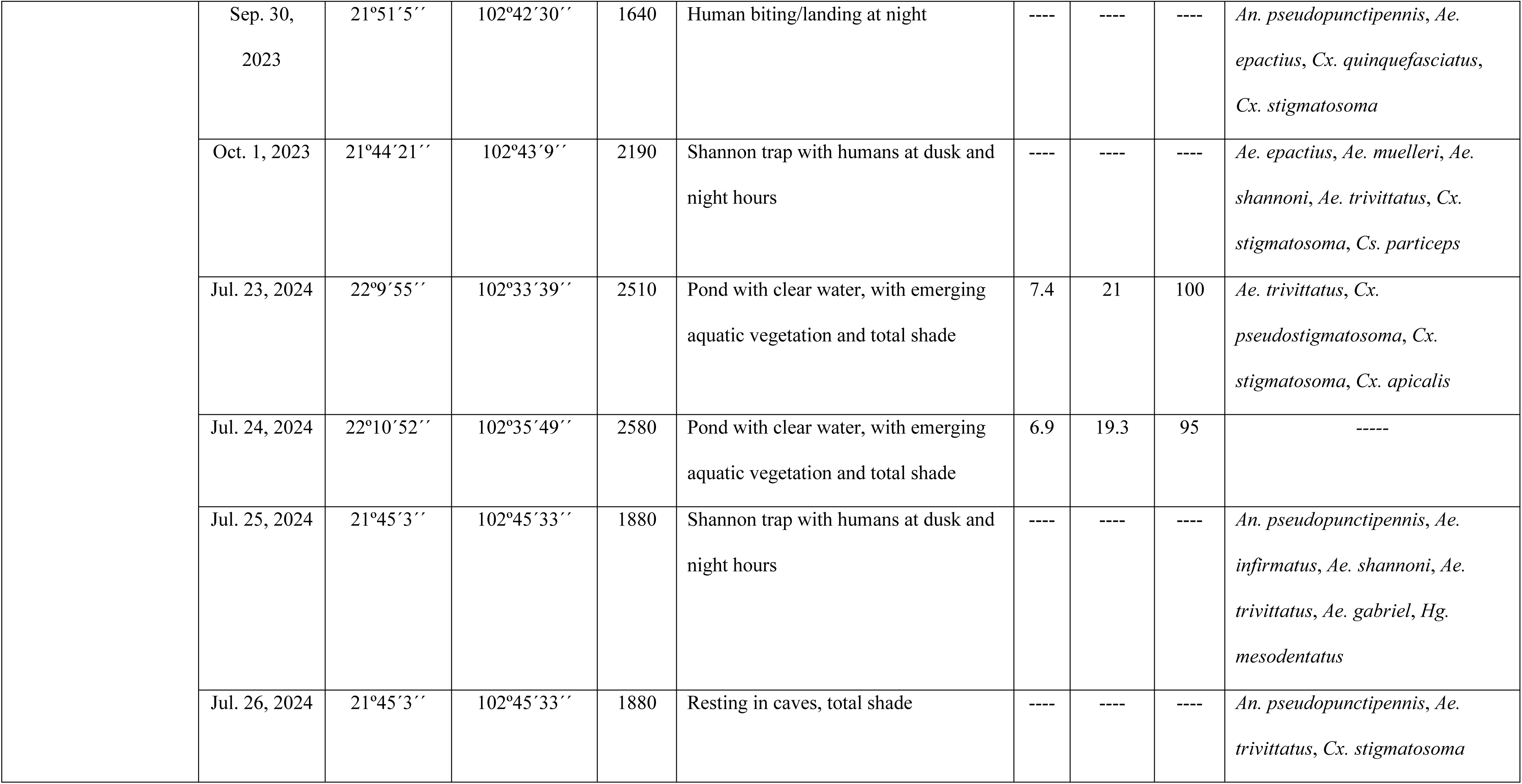
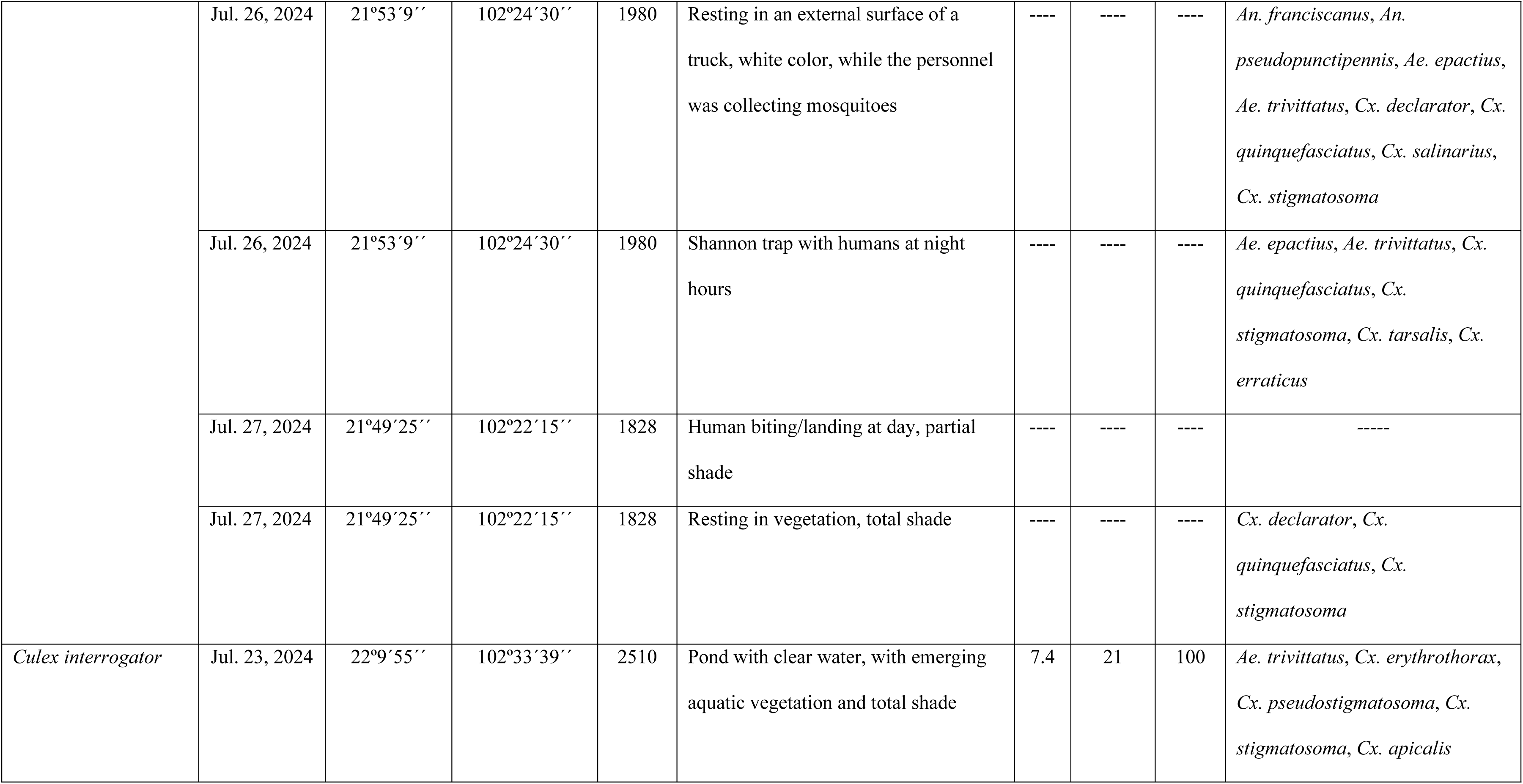
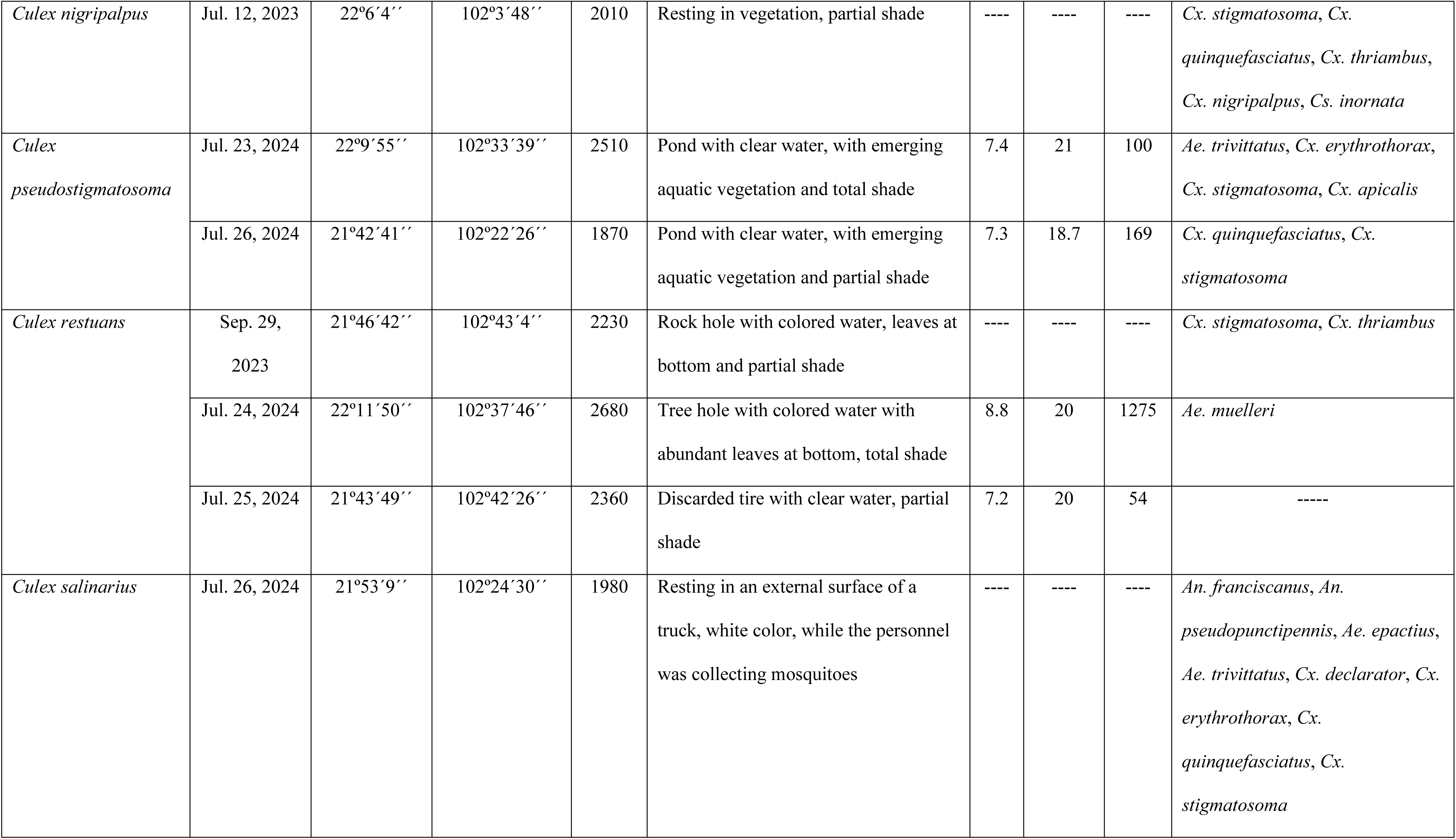
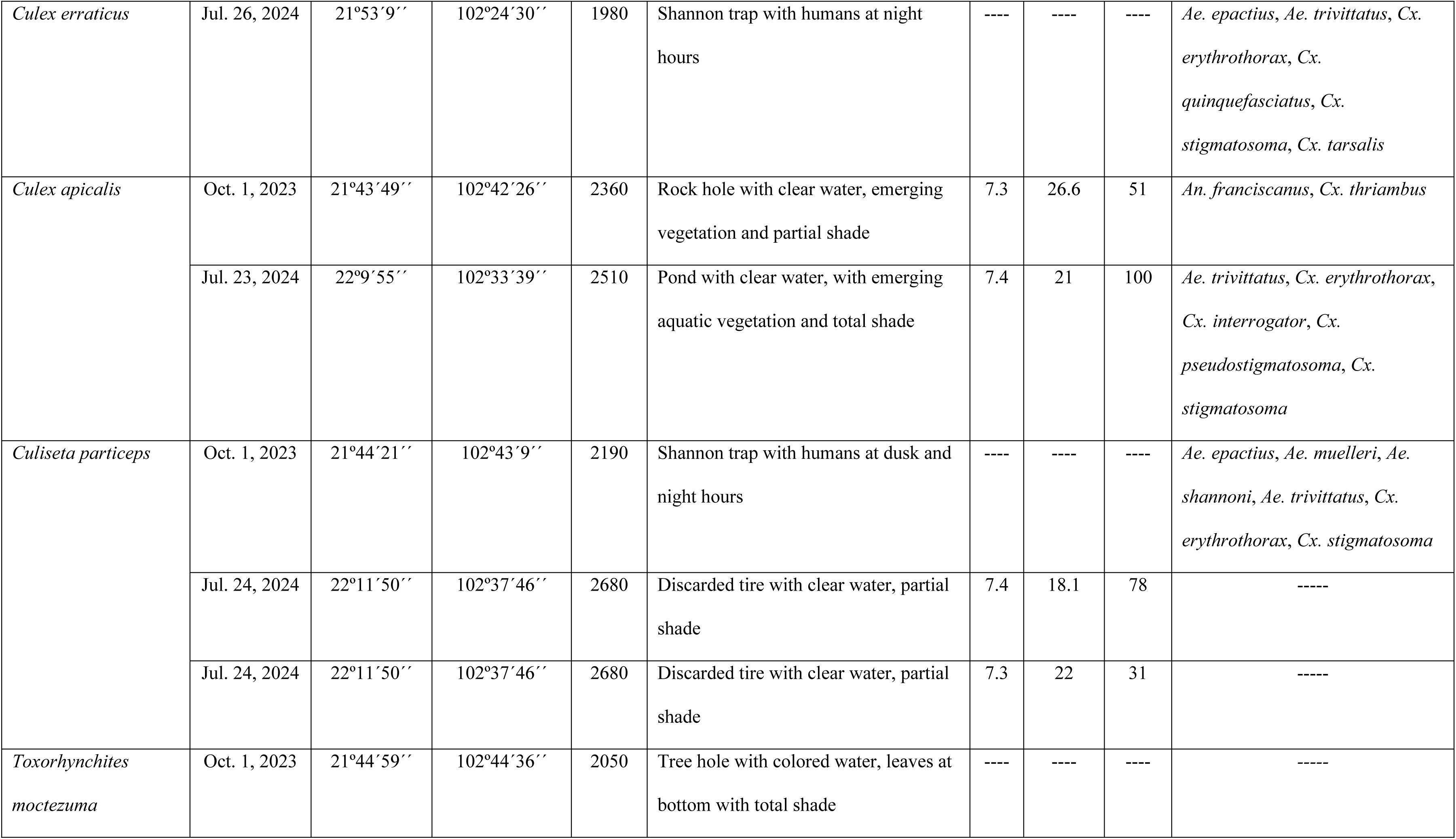
Collection records of the newly reported mosquito species found in Aguascalientes, Mexico. The position (geographic coordinate in scale of degrees, minutes and seconds); elevation (meters above sea level); type of aquatic habitat of the immature stages or environmental condition where adults were collected; environmental parameters of the aquatic habitat (pH, temperature and dissolved salts in scale of parts per million); and associated species for each collection.

### Review of Entomological Collection

The Collection of Arthropods of Medical Importance (CAIM) deposited in the Diagnostic and Epidemiologic Reference Institute (InDRE) in Mexico City, Mexico was examined for further additional records of Aguascalientes mosquitoes. The classification of the Culicidae family proposed by Wilkerson et al. [46] and the generic and subgeneric abbreviations of Culicidae names were considered.

### DNA Extraction and Molecular Barcoding

Genomic DNA from two legs (for- and hindlegs) per individual specimen, isolated using the modified hotshot technique [47], were placed directly into 50 µL of alkaline lysis buffer in micro vials and further sonicated in a water bath for 20 min. Micro vials were subsequently incubated in a thermocycler for 30 min at 94°C and cooled for 5 min. at 4°C, after which 50 µL of the neutralizing buffer was added to each vial. The cytochrome oxidase 1 (*COI*) barcode region was amplified using the primers LCO1490 (5′-GGTCAACAAATCATAAAGATATTGG-3′) and HCO2198 (5′-TAAACTTCAGGGTGACCAAAAAATCA-3′) [48]. Polymerase Chain Reactions (PCR) were performed using the MyTaq™ Mix (BIO-25041, Meridian Bioscience, OH, USA) and PCR conditions described by Ortega-Morales et al. [14]. The PCR products were electrophoresed in 1% agarose gel, and then fragments, with the expected amplicon length, were sequenced bidirectionally by Eurofins (Eurofins, Genomics, Louisville, KY, USA) using the traditional Sanger sequencing protocol [44].

All PCR products that reached the correct fragment size were subjected to bidirectional sequencing (Eurofins Genomics, KY, USA). The sequencing results were edited and analyzed using Chromas 2.6.6 (Technelysium, Pty, Ltd, Australia) and BioEdit version 7.1.1 and consensus sequence was generated using ClustalW [49]. Full details for each specimen and sequence information can be found at the National Center for Biotechnology Information (NCBI): *Anopheles pseudopunctipennis* (PZ832184, PZ832185); *Aedes epactius* (PZ832164, PZ832165); *Ae. shannoni* (PZ832166); *Ae. trivittatus* (PZ832167, PZ832168, PZ832169); *Ae. aegypti* (PZ832174); *Haemagogus mesodentatus* (PZ832170, PZ832171, PZ832172, PZ832173); *Culex quinquefasciatus* (PZ832176, PZ832177, PZ832178); *Cx. restuans* (PZ832179, PZ832180); C*x. stigmatosoma* (PZ832187, PZ832188, PZ832189, PZ832190, PZ832191, PZ832192, PZ832193, PZ832194, PZ832195, PZ832196); *Cx. tarsalis* (PZ832181, PZ832182, PZ832183); and *Culiseta inornata* (PZ832175). For certain species, we used complementary public *COI* barcode sequences publicly available in the Barcode of Life Data (BOLD): *An. pseudopunctipennis* (Mexico State, Mexico) (MOSQEMX022018G08, MOSQEMX022018E12, MOSQEMX022018F03, MOSQEMX022018F04); *Ae. burgeri* (Arizona, USA) (NEONTculicid2140, NEONTculicid2138, NEONTculicid2139); *Ae. nigromaculis* (Texas, USA) (NEONTculicid804, NEONTculicid2174, and NEONTculicid795); *Ae. gabriel* (Mexico State, Mexico) (QH1113EP6G05, QH1113EP6G03, QH1113EP6G10); *Ae. aegypti* (Mexico State, Mexico) (01210917 LBP1D8, 01210917 LBP1D7, QH1113EP6A12, QH1113EP6B01); *Psorophora signipennis* (Colorado, USA) (NEONTculicid803, NEONTculicid855, NEONTculicid801); *Cx. declarator* (Chiapas, Mexico) (VOLCAN032019D05, VOLCAN032019D04); *Cx. pseudostigmatosoma* (Mexico State, Mexico) (QH1113EP6D02, QH1113EP6D09, QH1113EP6D10, QH1113EP6D12); *Cx. salinarius* (Mexico State, Mexico) (MOSQEMX022018E04); *Cx. erraticus* (Tabasco, Mexico) (CUAC1414C-H01); *Cs. inornata* (Mexico State, Mexico) (07080917 TAP1A10, 07080917 TAP1A8); and *Cs. particeps* (Mexico State, Mexico) (MOSQEMX022018H03, MOSQEMX022018H06, 07080917 TAP1A7, MOSQEMX022018H05). The dataset was analyzed in MEGA v.6 [50]. The Maximum Likelyhood (ML) analysis was performed using the Kimura 2-Parameter distance metric to determinate their distribution pattern, and the tree was rooted to *An. gambiae* Giles (Nigeria), (GM84-NGA). The tree robustness was measured by the bootstrap approach using 1,000 pseudoreplicates.

### Bloodmeal Metabarcoding Methods

DNA from the abdomens of 64 engorged mosquitoes were extracted using the Thermo Scientific™ KingFisher™ Flex Purification System and MagMAX™ Core Nucleic Acid Purification Kit (Thermo Fisher Scientific, Waltham, MA, USA) following previously published protocols [51]. All DNA extracted from samples were tested by PCR using 12S rRNA mitochondrial locus primers with dual identical barcode tags [52–54]. Primers were 12S-6F CAAACTGGGATTAGATACC and 12S-13R AGGAGGGTGACGGGCGGT [55] ∼145 bp in length, with identical 10 bp barcode tags [56]. PCR was performed in duplicate for each sample with separate barcode tags for each replicate. Of the 64 samples tested, 23 samples amplified in replicate and thus were further processed. These PCR products were pooled and subject to library preparation with xGenTM ssDNA & Low Input DNA Library Prep kit from Integrated DNA technologies (Coralville, IA, USA) and sequencing was run on the Illumina NextSeq2000 platform at the Texas Institute for Genome Sciences (https://genomics.tamu.edu).

Sequences were demultiplexed on dual identical barcode tags with tag trimming using Cutadapt 5.0 [57] on a minimum length of 100 bp. Subsequent binning and filtering was done using Qiime2 - primers were trimmed, sequences were merged and denoised at minimum length of 100 bp and truncation at 145 bp both forward and reverse [58]. To identify hosts, sequences were matched to taxa using NCBI BLAST (National Center for Biotechnology Information, US National Library of Medicine) with MegaBLAST with the GenBank database (https://www.ncbi.nlm.nih.gov/genbank/) [59]. Only sequences (Amplicon Sequence Variants, ASVs) that had more than 100 reads across the study were retained. Identity match of 99% or higher to the GenBank sequences was required for species-level identification; lower percent matches were binned into family or order. If multiple species showed equal query cover, e-value and identity match, geographic distribution was taken into consideration to rule out species that do not occur in the region. Genbank matches used the identifier *Bubulucus ibis* for the cattle egert. Current nomenclature of *Adrea ibis* is being used in this study. The lowest common taxonomic rank that fits the geographic distribution of the host matches was accepted (Columbidae, Passeriformes). Information for host distribution was obtained from GBIF (https://www.gbif.org/). If information for birds was unavailable or limited on GBIF, Avibase (https://avibase.bsc-eoc.org) was used as a secondary reference.

## Results

### Mosquito Identification

Overall, 5,227 specimens from 111 collections were examined: 357 fourth-instar larvae, 199 larval exuviae, 373 pupal exuviae, 33 pupae, 3,000 adult females, 1,264 adult males, and 1 dissected male genitalia. The mosquito fauna of Aguascalientes consists of 34 species representing the subfamilies Anophelinae and Culicinae, four tribes of the subfamily Culicinae, seven genera, and 14 subgenera (Table 3). One tribe (Toxorhynchitini), three genera (*Haemagogus*, *Psorophora*, and *Toxorhynchites*), nine subgenera (*Aedimorphus*, *Lewnielsenius*, *Ochlerotatus*, *Protomacleaya*, *Haemagogus*, *Grabhamia*, *Melanoconion*, *Neoculex*, and *Megarhina*), and 22 species (*Aedes vexans*, *Ae. muelleri*, *Ae. infirmatus*, *Ae. nigromaculis*, *Ae. shannoni*, *Ae. trivittatus*, *Ae. burgeri*, *Ae. gabriel*, *Haemagogus mesodentatus*, *Psorophora signipennis*, *Culex bidens*, *Cx. declarator*, *Cx. interrogator*, *Cx. erythrothorax*, *Cx. nigripalpus*, *Cx. pseudostigmatosoma*, *Cx. restuans*, *Cx. salinarius*, *Cx. erraticus*, *Cx. apicalis*, *Culiseta particeps*, and *Toxorhynchites moctezuma*) are new records (Table 3).

**Table 3.**
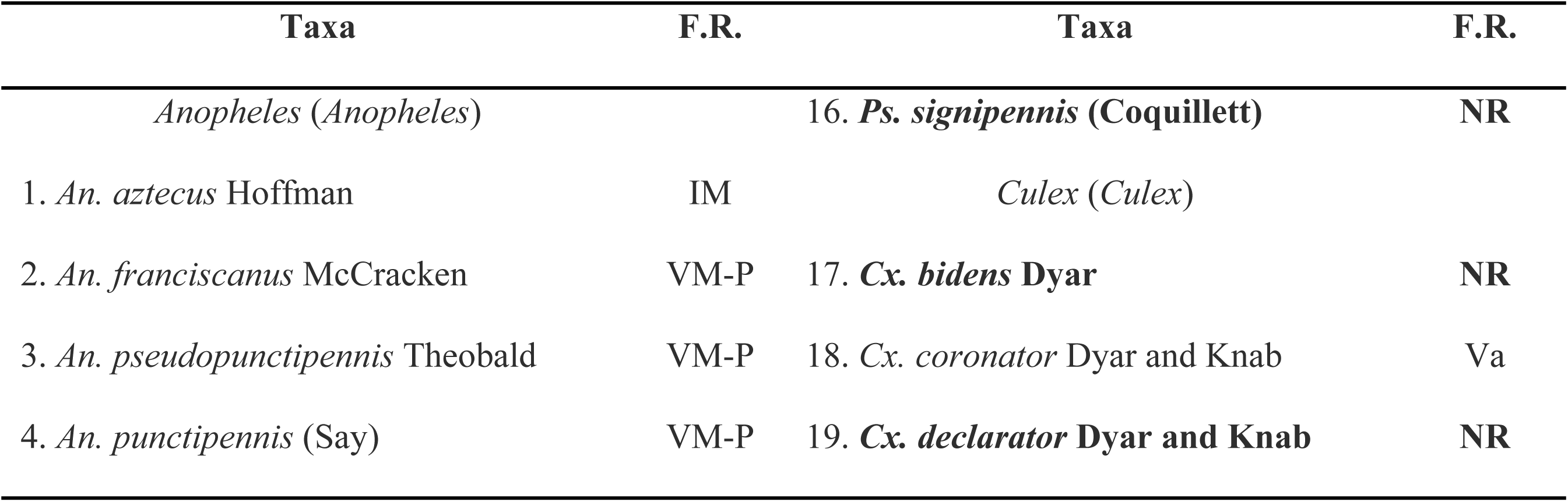
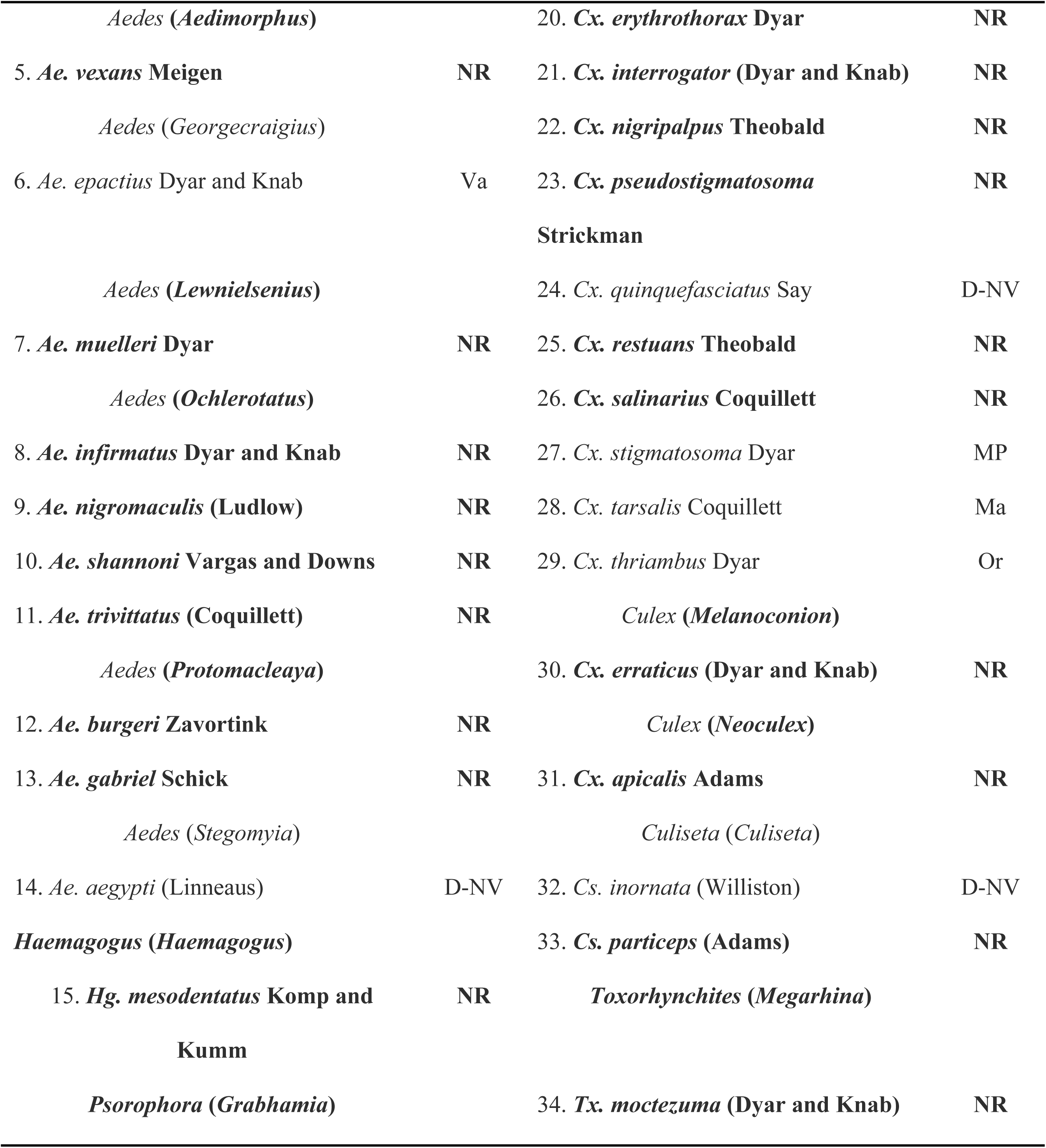
Mosquito species from Aguascalientes. IM: [**60]; VM-P: [3]; Va: [61]; D-NV: [62]; MP: [63]; Ma: [64]; Or: [65]. NR: New Record.**

The species accumulation curve shows a rapid initial increase in species discovery, reaching 25 species at sample 70 and 30 species at sample 100 (Fig. 2). The curve reaches a brief plateau during the final sampling interval (samples 100–111), indicating high sampling efficiency.

**Figure 2.**
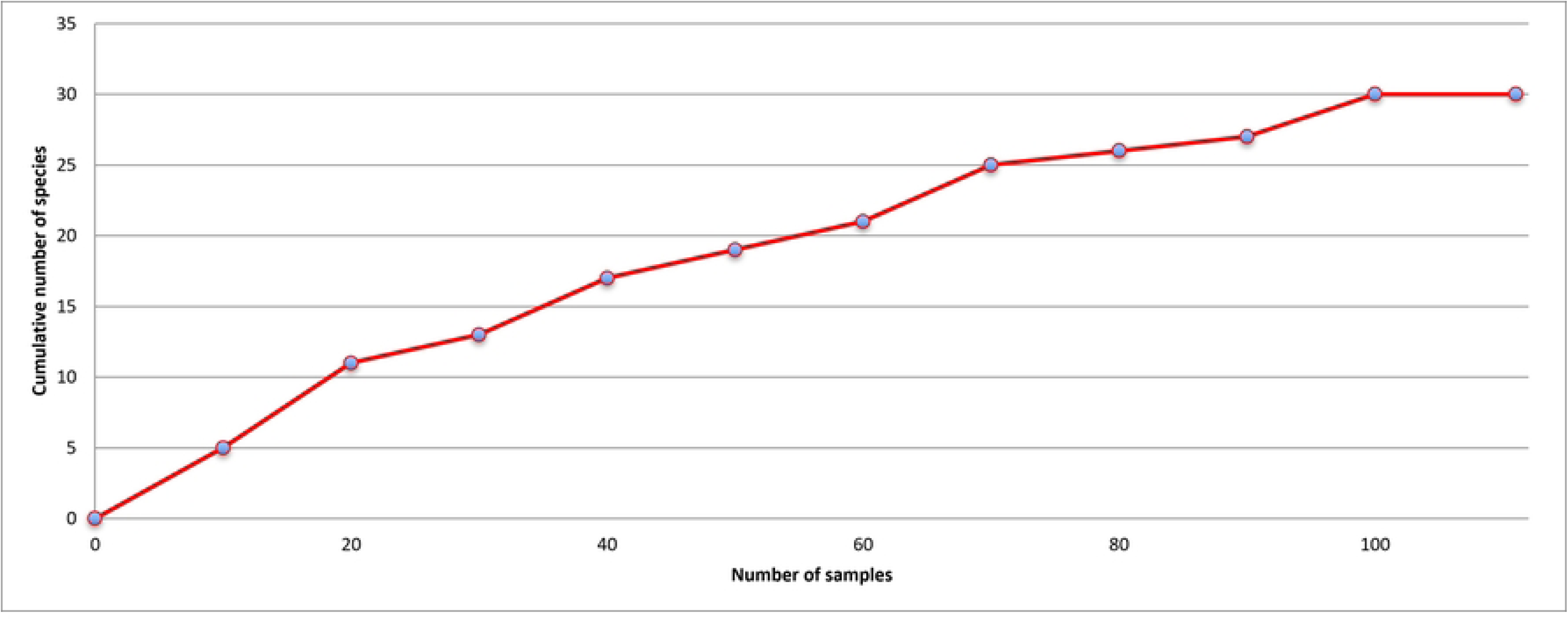
Species accumulation curve for 30 mosquito species (111 collections) collected in Aguascalientes during 2023–2024.

### New Records

#### Genus *Aedes*

Eight out 10 *Aedes* spp. of Aguascalientes are new records: *Ae. vexans*, *Ae. muelleri*, *Ae. infirmatus*, *Ae. nigromaculis*, *Ae. shannoni*, *Ae. trivittatus*, *Ae. burgeri*, and *Ae. gabriel*. The subgenera *Aedimorphus*, *Lewnielsenius*, *Ochlerotatus*, and *Protomacleaya* are also new records. Immature stages of *Ae. vexans* were collected once from a pond during the rainy season; other new record is *Ae. muelleri*, which is uncommon species in Mexico. In the “Sierra Madre Occidental” region, it can be found during the rainy season. Immature stages of *Ae. muelleri* were collected from six tree holes filled with rainwater, while adults were collected resting inside one of those tree holes. Host-seeking females were collected using Shannon traps baited with protected human volunteers, although some of these females only landed on clothing without biting intention. In the Central Plateau region, immature stages of *Ae. muelleri* were also collected from a discarded tire. Although the subgenus *Ochlerotatus* is common in Mexico, the documentation of taxa in this subgenus in Aguascalientes is a new record. Host-seeking females of *Ae. infirmatus* were collected once using Shannon traps at dusk hours and some of them land on the collector personnel with biting intention: *Ae. nigromaculis* is another uncommon species in Mexico, and host-seeking females of this species were collected landing on the collector personnel at day hours in the “Sierra Madre Occidental” and Neo-Volcanic Axis regions, one site by each region.

*Aedes shannoni* is a Mexican endemic species which distribution is restricted to the Mexico valley in the south-middle of Mexico. Aguascalientes is the northernmost distributional limit for this species; immature stages were collected form a rock hole and one discarded tire, while host-seeking females were collected using Shannon traps and landing on humans.

*Aedes trivittatus* is one of the most common species in Mexico but in Aguascalientes is a new record, immature stages were collected form five ponds at ground level with partial and total shade across Aguascalientes. Adults were collected resting in vegetation and caves with total shade, while host-seeking females were collected from all regions, landing on humans with biting intention at day and night hours, resting on a truck, and using Shannon traps at dusk and night hours.

Finally, the subgenus *Protomacleaya* and the species *Ae. burgeri* and *Ae. gabriel* are new records. Immature stages of *Ae. burgeri* were collected from one tree hole in a live oak tree (*Quercus* sp.) at 1 m. from ground level while host-seeking females of *Ae. gabriel* were collected in the “Sierra Madre Occidental” region landing on humans with biting intention and using Shannon traps at dusk and night hours.

#### Genus *Haemagogus*

The genus *Haemagogus*, subgenus *Haemagogus* and the species *Hg. mesodentatus* are new records. In the Sierra Madre Occidental region, host -seeking females were collected landing on humans during day hours and using Shannon traps at dusk and during night hours.

#### Genus *Psorophora*

In Mexico, the genus *Psorophora* is a common group of mosquitoes during the rainy season; however, this is new record of this genus, subgenus *Grabhamia*, and the species *Ps. signipennis* in Aguascalientes. Immature stages of this species were collected once from one pond during the rainy season in the Central Plateau region.

#### Genus *Culex*

The genus *Culex* is the most diverse group of mosquitoes in Aguascalientes, including 15 species, of which only five had previously been reported. The other ten species; *Culex bidens*, *Cx. declarator*, *Cx. erythrothorax*, *Cx. interrogator*, *Cx. nigripalpus*, *Cx. pseudostigmatosoma*, *Cx. restuans*, *Cx. salinarius*, *Cx. erraticus*, and *Cx. apicalis* are reported new records. Immature stages of *Cx. bidens* were collected from one site of the Central Plateau region, in a pond with total shade; immature stages of *Cx. declarator* were collected from a rock hole in a single site in the Central Plateau region, and adults were collected resting in vegetation and resting in a truck; *Cx. erythrothorax* is the most common species within the genus *Culex* in Aguascalientes, is new record. Immature stages were collected from ponds with emerging vegetation in the Central Plateau region, while adults were collected resting in vegetation, caves and intra-domiciliary. Host-seeking females were collected landing on humans at day and night hours, resting in a truck, and using Shannon traps at night hours, but this host-seeking females only landed on clothes without biting intention. One single larvae of *Cx. interrogator* was collected from a pond with emerging vegetation in the “Sierra Madre Occidental” region. Adults of *Cx. nigripalpus* were collected once resting in vegetation with partial shade in the Central Plateau region; *Culex pseudostigmatosoma* is one uncommon tropical species in Mexico, but this species is apparently common in the Mexico valley and adjacent semi-arid areas. Immature stages were collected from two ponds at ground level with abundant emerging vegetation in the Central Plateau region, where this species reach its northernmost distributional limit. Immature stages of *Cx. restuans* were collected from a variety of aquatic habitats serving as breeding sites such as rock and tree holes, and discarded tires. Adults of *Cx. salinarius* were collected resting in a truck. The subgenus *Melanoconion* and the species *Cx. erraticus* are new records. Host-seeking females were collected using Shannon traps with humans at night hours, landing on clothes without biting intention. Finally, the subgenus *Neoculex* and the species *Cx. apicalis* are new records. Immature stages were collected from one rock hole and one pond with clear water in the “Sierra Madre Occidental” region.

#### Genus *Culiseta*

*Culiseta particeps* is the most common species within the genus *Culiseta* in Mexico, however, this is new record for Aguascalientes. In the “Sierra Madre Occidental” region, immature stages of *Cs. particeps* were collected from discarded tires with partial shade, while host-seeking females were collected using Shannon traps with humans at night hours, landing on clothes without biting intention.

#### Genus *Toxorhynchites*

The tribe Toxorhynchitini, genus *Toxorhynchites*, subgenus *Megarhina* and *Tx. moctezuma* are new records. One single larval exuviae of *Tx. moctezuma* was found from one oak tree hole with colored water in the “Sierra Madre Occidental region”. Interestingly, no additional specimens were found on this site.

### DNA barcoding and phylogenetic analysis

Overall, we analyzed 78 DNA barcodes for 20 species, covering 58.8% of total species of Aguascalientes. Phylogenetic relationships were reconstructed by Bayesian inference (Fig. 3). We could not obtain DNA barcodes for five species as they were collected as immature stages and previously mounted on microscope slides for morphological identification: *An. franciscanus*, *Cx. bidens*, *Cx. interrogator*, *Cx. apicalis*, and *Tx. moctezuma*. A further five species were processed but failed to amplify DNA: *Ae. vexans*, *Ae. infirmatus*, *Cx. erythrothorax*, *Cx. nigripalpus*, and *Cx. thriambus*.

**Figure 3.**
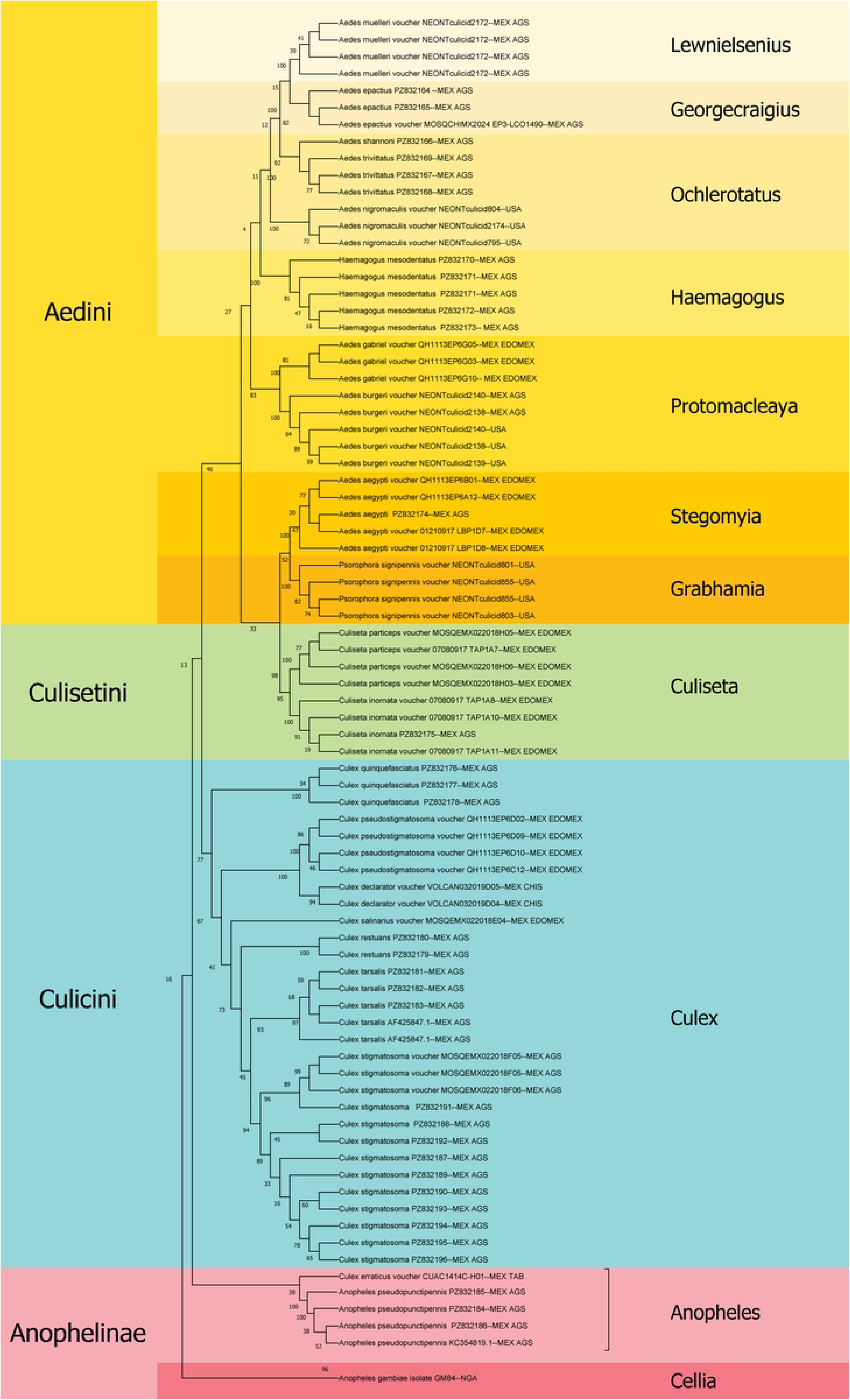
Maximum likelihood tree base on the Kimura 2-parameter of the *COI* DNA barcodes (658 bp) for 20 mosquito species of Aguascalientes.

As a rule, specimens of the same species clustered together, although there was a deep split in *Cx. stigmatosoma* (MOSQUEMX022018F05, MT108648.1, MT108615.1). Within the tribe Aedini, in the genus *Aedes,* there are two strong groups of monophyletic species with boostrap values over 80%. The subgenus *Ochlerotatus*: *Ae. shannoni* (JX259638.1), *Ae. trivittatus* (MG242524.1, JX259690.1), and *Ae. nigromaculis* (NEONTculicid804, NEONTculicid2174, NEONTculicid795). *Aedes shannoni* and *Ae. trivittatus* are placed at the same clade. In larval stage, *Ae. shannoni* showed affinities to the Scapularis Group [14] which *Ae. trivittatus* belongs. The subgenus *Protomacleaya*: *Ae. gabriel* (QH1113EP6G05, QH1113EP6G03, QH1113EP6G10) and *Ae. burgeri* (NEONTculicid2140, NEONTculicid2138, NEONTculicid2139); and species in different subgenus but placed in the same clade as *Ae. muelleri* (NEONTculicid2172) in the subgenus *Lewnielsenius* and *Ae*. *epactius* (PP549542.1, MT108613.1, and MOSQCHIMX2024 EP3-LCO1490) in the subgenus *Georgecraigius*. Other members of the tribe Aedini were *Hg. mesodentatus* (MT108553.1), *Ae. aegypti* (QH1113EP6B01, QH1113EP6A12, KY022526.1, 01210917 LBP1D7, 01210917 LBP1D8), and *Ps. signipennis* (NEONTculicid801, NEONTculicid855, NEONTculicid803), with a single clade respectively. Within the tribe Culicini, genus *Culex* and subgenus *Culex*, formed monophyletic group >60% *Cx. quinquefasciatus* (MK714012.1, MH463059.1), *Cx. pseudostigmatosoma* (QH1113EP6D02, QH1113EP6D09, QH1113EP6D10, QH1113EP6C12), *Cx. declarator* (VOLCAN032019D05, VOLCAN032019D04), *Cx. salinarius* (MOSQEMX022018E04), *Cx. restuans* (PP558251.1, PP558247.1), *Cx. tarsalis* (AF425847.1), and *Cx. stigmatosoma* (MOSQEMX022018F05, MOSQEMX022018F06, MT108648.1, MT108673.1, MT108615.1); *Cx. erraticus* (CUAC1414C-H01) is the only species in the subgenus *Melanoconion* included in the phylogenetic relationship. Finally, the genus *Anopheles*, subgenus *Anopheles*: *An. pseudopunctipennis* (MT034039.1, KC04039.1, KC354819.1) and the genus *Culiseta*, subgenus *Culiseta*: *Cs. particeps* (MOSQEMX022018H05, 07080917 TAP1A7, MOSQEMX022018H06, MOSQEMX022018H03), and *Cs. inornata* (07080917 TAP1A8, 07080917 TAP1A10, PP558229.1, 07080917 TAP1A11) formed singled and separated clades, respectively.

### Host Feeding Patterns

Bloodmeal host information was obtained from 23 of 64 (36%) mosquitoes of the following species: *Ae. epactius* (1), *An. pseudopunctipennis* (1), *Cs. inornata* (1), *Cx. quinquefasciatus* (6) and *Cx. stigmatosoma* (14) (Table 4). The resolution of analysis afforded detection of seven species of vertebrates in addition to one family and one order for which species could not be determined. Mammals accounted for four species and one order, whereas birds accounted for three species and one family (Table 5). Birds were the most frequent hosts, accounting for 46 of the 67 bloodmeals hosts detected. *Ardea ibis* L. (cattle egret) was the most frequent host (n=18) followed by *Zenaida asiatica* L. (white-winged dove) (n=12). In both, *Cx. quinquefasciatus* (60% of 15 bloodmeals) and *Cx. stigmatosoma* (68% of 41 bloodmeals) most bloodmeals detected were again avian hosts.

**Table 4:** Table showing species of mosquitoes and the number of vertebrate hosts species that were identified from each species.

| Mosquito species | Number in study | Number with host/s identified | Number of host species detected in bloodmeals |  |  |  |  |  |
| --- | --- | --- | --- | --- | --- | --- | --- | --- |
|  |  |  | 1 | 2 | 3 | 4 | 5 | 6 |
| *Ae. epactius* | 1 | 1 |  |  |  | 1 |  |  |
| *An. pseudopunctipennis* | 1 | 1 | 1 |  |  |  |  |  |
| *Cs. inornata* | 2 | 1 |  |  |  | 1 |  |  |
| *Cx. quinquefasciatus* | 21 | 6 | 1 | 2 | 2 |  |  | 1 |
| *Cx. stigmatosoma* | 36 | 14 | 4 | 1 | 3 | 4 | 2 |  |

**Table 5.**
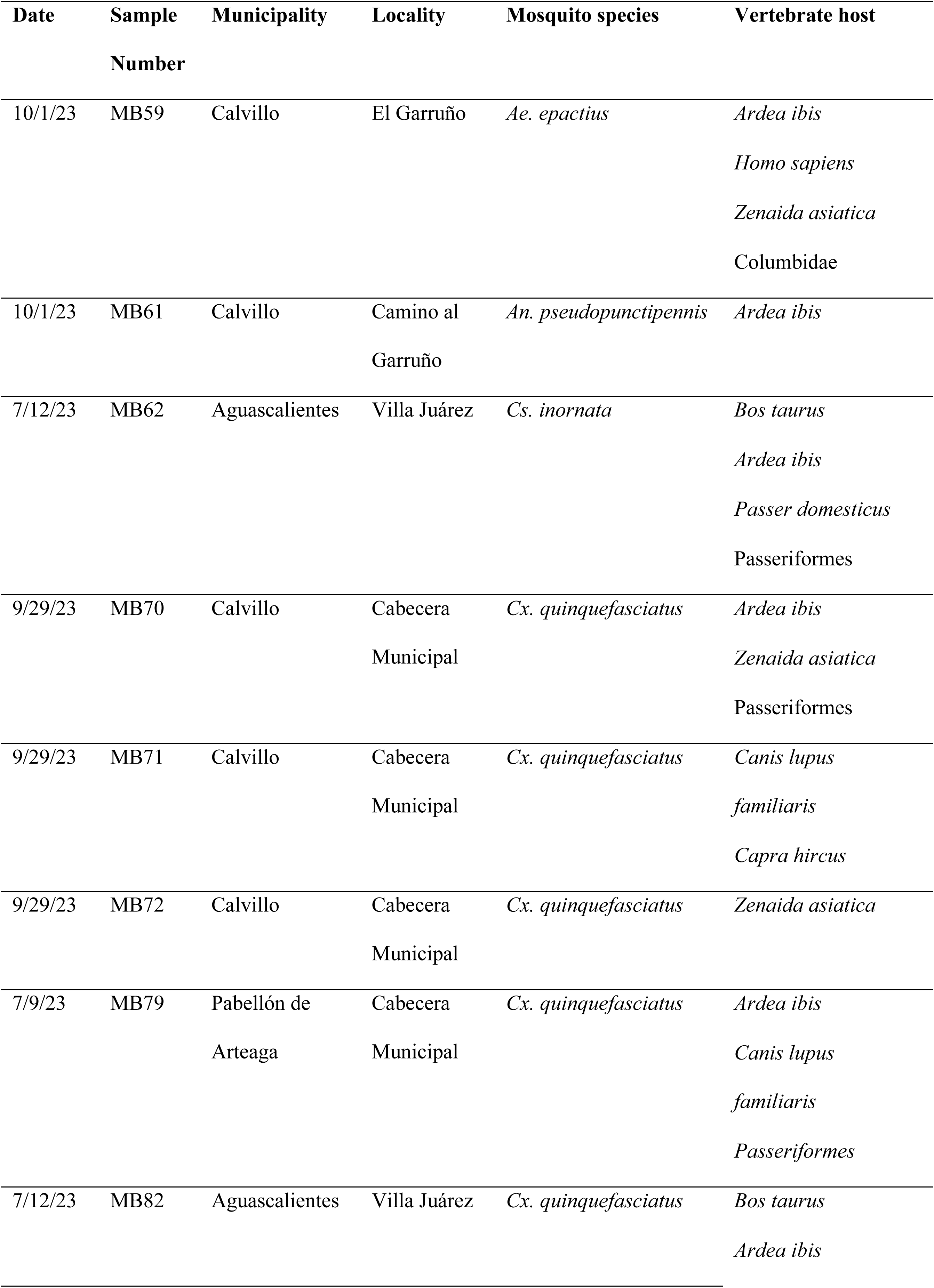
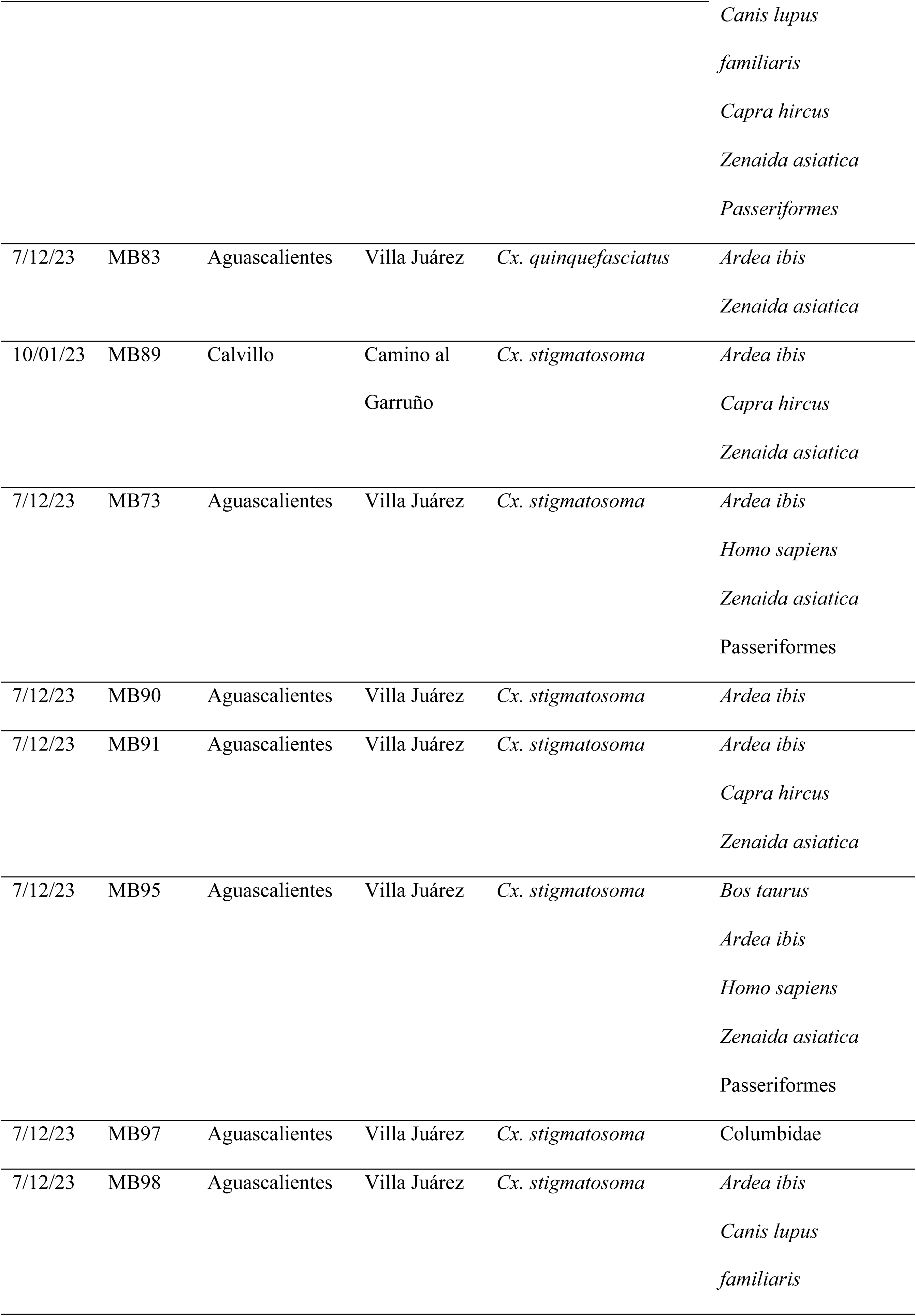
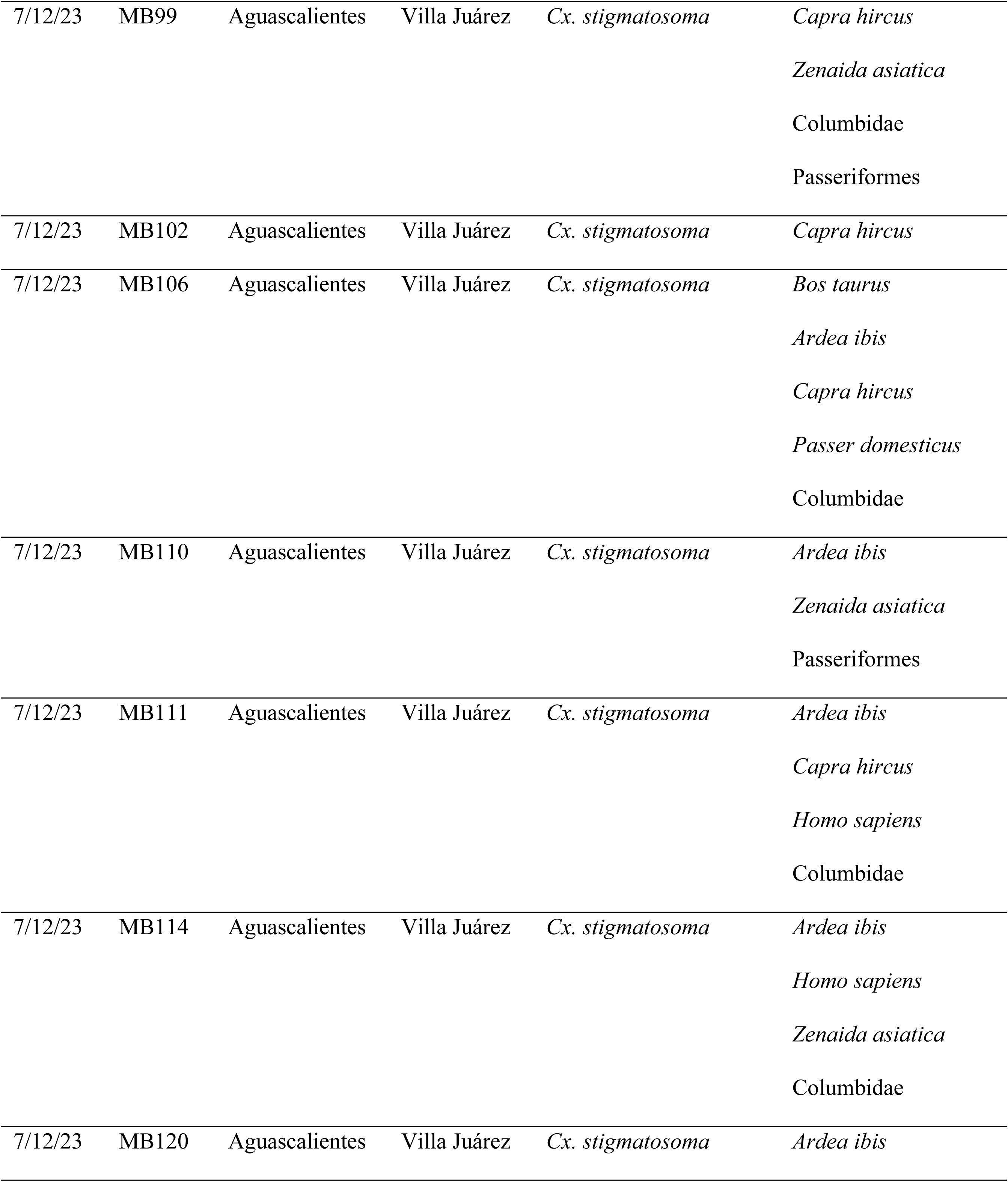
Summary of vertebrate host species identified from mosquitoes collected from Calvillo and Aguascalientes municipalities.

Most mosquitoes had evidence of having fed upon more than one vertebrate species (19/23 = 83%) (Table 4–5, Figure 4). The maximum number of host species detected from an individual mosquito was six; this occurred in *Cx. quinquefasciatus* in which *Bos taurus* L. (cow), *Ardea ibis* (cattle egret), *Canis lupus familiaris* L. (dog), *Capra hircus* L. (goat), *Z. asiatica* (white-winged dove), and a bird of the order Passeriformes were co-detected. The single host meals were *A. ibis* (3), *C. hircus* (1), *Z. asiatica* (1), and family-level host, Columbidae (1). Human hosts were detected in five instances but never as the sole host bloodmeals; four human meals were found in *Cx. stigmatosoma* (Figure 4).

**Figure 4.**
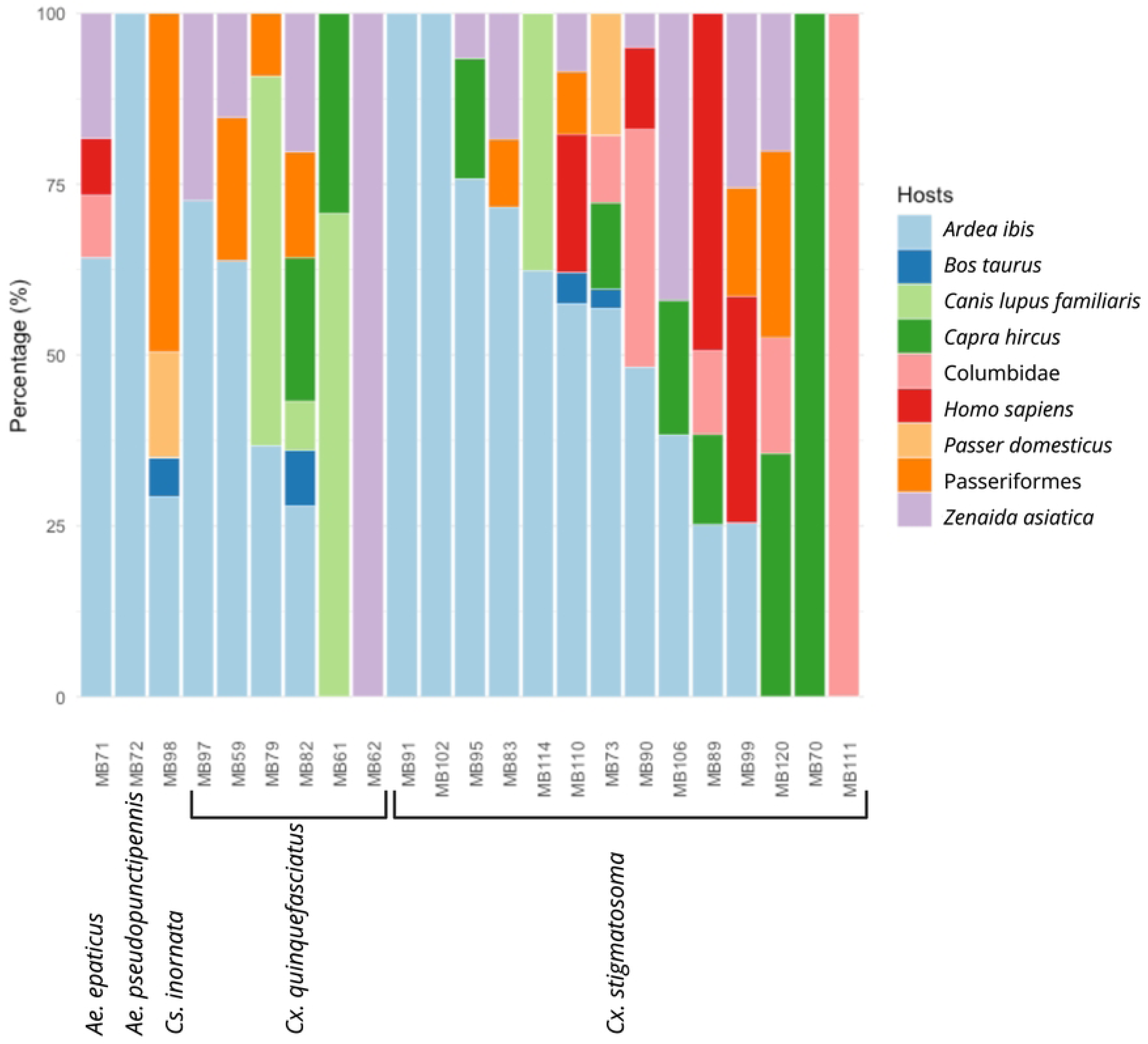
Stacked bar graph shows host use in individual mosquitoes. Reads per host are represented as percent values. Figure was generated in R ggplot2 (Wickhan H (2016). *ggplot2*: Elegant Graphics for Data Analysis. Springer–Verlag New York. ISBN 978-3-319-24227-4). Labels were adjusted for aesthetics with BioRender.com

## Discussion

### Mosquito richness of Aguascalientes and Mexico

Here, 22 species are new records of mosquitoes for Aguascalientes; thus, there are 33 species documented to occur in Aguascalientes. Aguascalientes ranks eleventh for species richness out of 13 Mexican states, which has been systematically inventoried. With the removal of *Aedes idahoensis* of Mexico, there are 255 species of mosquitoes in Mexico, overall.

### Blood feeding patterns

The main hosts identified in analysis of bloodmeals suggest birds, mainly the cattle egret, columbids and passerines as predominant hosts. Aside from birds, livestock, dogs, and humans were mammalian hosts. Only 32% of the bloodmeals were single host meals. Of the 19 multiple host bloodmeals, three (two *Cx. quinquefasciatus*, one *Cx. stigmatosoma*) were all bird hosts and one was all mammal hosts. The remaining 13 (68% of mixed bloodmeals detected) included a mix of bird and mammal bloodmeals. This observation, with other studies on larger sample sizes of *Cx. quniquefasciatus* bloodmeals from Chiapas, Tamaulipas, and Texas confirm that these mosquitoes freely fed across vertebrate classes. Of the medically relevant species known to vector pathogens (Table 6), two *Cx. quniquefasciatus* mosquitoes showed avian and mammal mixed bloodmeals and two mixed bird and one mixed mammal meal. *Culiseta inornata* showed a mixed bird bloodmeal. This mixture of meals points toward a concrete pathway of transmission of mosquito-borne viruses and protozoans and their maintenance in enzootic cycles. Our observation of six different host species within a single individual underscores the complex ecological associations among wildlife and domestic animals that sustain the mosquitoes in nature.

**Table 6.**
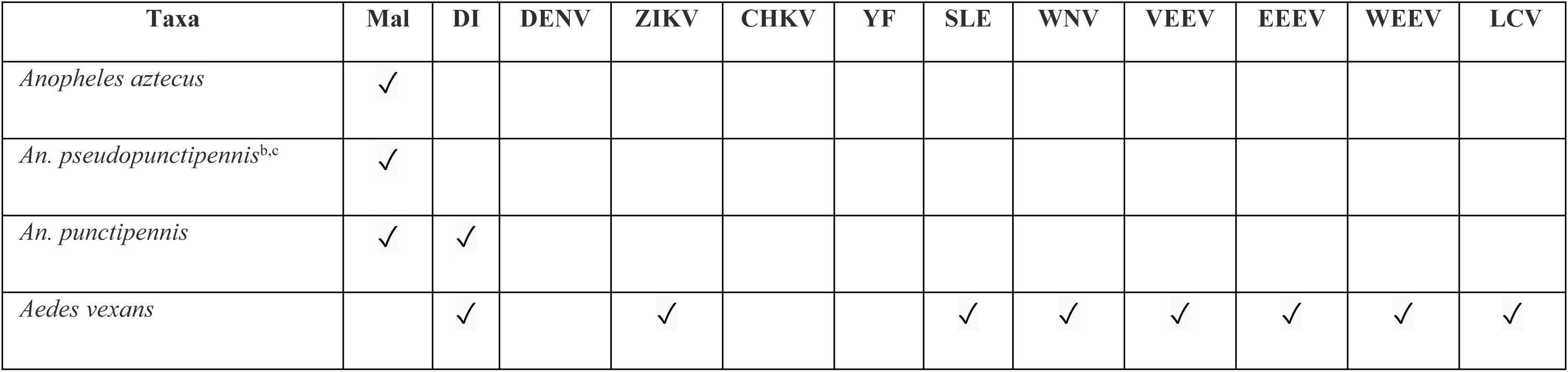
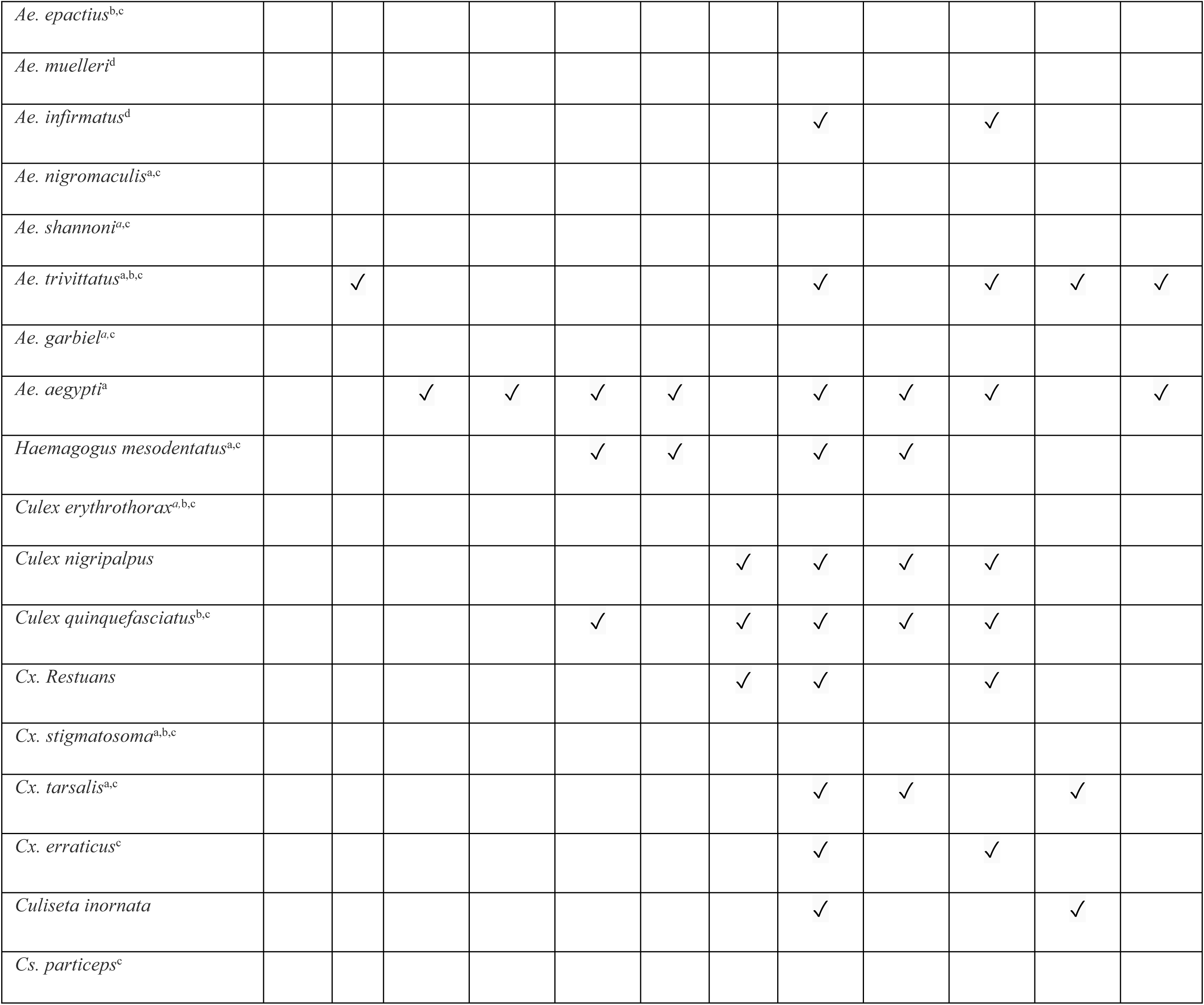
Medically important mosquitos of Aguascalientes as they may serve as vectors of parasite and viral pathogens. Mal: Malaria. DI: *Dirofilaria immitis*. DENV: dengue virus. ZIKV: Zika virus. CHIKV: chikungunya virus. YF: yellow fever virus. SLE: St. Louis encephalitis virus. WNV: West Nile virus. VEEV: Venezuelan equine encephalitis virus. EEEV: eastern equine encephalitis virus. WEEV: western equine encephalitis virus. LCV: La Crosse virus. ^a^Host-seeking mosquitoes caught using human landing collections at day light hours. ^b^Host-seeking mosquitoes caught using human landing collections at night hours. ^c^Host-seeking mosquitoes caught using Shannon traps at dusk and night hours.

### Species from adjacent regions that may occur in Aguascalientes

Some species of mosquitoes that have not yet been reported from Aguascalientes occur in adjacent Jalisco and Zacatecas; hence, the occurrence in Aguascalientes of those species could not be ruled out. These are two species for Zacatecas: *Anopheles judithae* Zavortink and *Orthopodomyia kummi* Edwards and two for Jalisco: *Aedes niveoscutum* Zavortink, and *Cx. restrictor* Dyar and Knab [66–68]. Species reported in non-adjacent Queretaro, but a close state that shares physiographic similarities with Aguascalientes, are [7]: *An. apicimacula* Dyar and Knab, *An. eiseni* Coquillett, *An. albimanus* Wiedemann, *Ae. scapularis* (Rondani), *Cx. peccator* Dyar and Knab, *Cx. arizonensis* Bohart, *Cx. lactator* Dyar and Knab, *Lutzia bigoti* (Bellardi), and *Culiseta dugesi* Dyar and Knab. Those species in Mexico State [9]: *Ae. ramirezi* Vargas and Downs, *Ae. zoosophus* Dyar and Knab, *Cs. incidens* (Thomson), and *Uranotaenia geometrica* Theobald. And those species in Hidalgo [11–13]: *Psorophora ferox* (von Humboldt), *Cx. corniger* Theobald, *Uranotaenia geometrica* Theobald, and *Ur. sapphirina* (Osten Sacken).

### Reported species of Aguascalientes which could not be verified in our collections

Three species were previously reported from Aguascalientes but those were not found in our collections: *Anopheles aztecus*, *An. punctipennis*, and *Culex coronator*. *Anopheles aztecus* is a Mexican endemic species that have been reported in some states of the Mexico Valley, as well in Durango and Veracruz. Recent collection records for this species are scarce, being one of the uncommon *Anopheles* species in Mexico. By other hand, *An. punctipennis* is a common species with many Mexican recent collection records. This species is particularly common in arid and semi-arid areas; however, we did not collect this species during the current study in Aguascalientes. *Culex coronator* is one of the most common mosquito species in Mexico, however, this species was not found in our collections. In the CAIM, we found 24 records belonging to *An. punctipennis* collected mostly between 1941–1955 by A. Díaz-Nájera, A. Martínez-Palacios, and V. Molina-Valente in the counties of Aguascalientes and Calvillo. Two records for *Cx. coronator* collected in 1941 and 2010 by A. Martínez-Palacios, and E. Islas Ojeda in the counties of Aguascalientes and Calvillo, respectively were also noted. The species accumulation curve suggests that additional sampling during this study would have detected additional species, as we had not yet reached an asymptote with no new species being discovered with additional sampling effort.

### The Removal of *Aedes idahoensis* (Theobald) from Mexico

*Aedes* (*Ochlerotatus*) *spencerii* was originally described by Theobald [69] as *Culex spencerii* from four adult females collected by W. I. Spencer in Manitoba, Canada. Original illustrations of wing, abdominal segment in dorsal aspect, and base of antenna of female were provided from Theobald [69]. Lectotype material was selected by Belkin from type locality in Stony Mountain, Manitoba, Canada and deposited in the British Museum of Natural History (BMNH) [70]. *Aedes spencerii* is distributed in the Interior Plains region of Canada in the provinces of Alberta, British Columbia, Manitoba, Saskatchewan, Ontario and Yukon Territory, and Unites States in the states of Illinois, Iowa, Kansas, Michigan, Minnesota, Montana, Nebraska, New York, North Dakota, Ohio, Oklahoma, South Dakota, Wisconsin, and Wyoming [71–73]. *Aedes* (*Ochlerotatus*) *idahoensis* was originally described by Theobald [74] as *Grabhamia idahoensis* variety (= subspecies) of *Ae. spencerii* from an unspecified number of specimens collected by J. M. Aldrich in Market Lake, Idaho, United States. Two syntype females are deposited in the BMNH [75]. *Aedes idahoensis* is distributed in southern Canada in the province of British Columbia and Northwestern United States in the States of Colorado, Idaho, Montana, Nebraska, Nevada, North Dakota, Oregon, South Dakota, Utah, Washington, and Wyoming [76–77]. Theobald [74] stated *Ae. idahoensis* differs from *Ae. spencerii* in abdominal scale ornamentation which described as “Abdominal basal bands almost white, broad apical bands yellow-white to white, very narrow, almost obliterated on some segments; the abdomen not pale at the sides, so that only broad basal and narrow apical pale areas are shown, the two apical segments are mostly pale scaled, and there are scattered pale scales on the others” while in the typical *Ae. spencerii*, Theobald [69] indicate “Abdomen covered with creamy and white scales, with large, black-scaled patches on each side of the middle line”. *Aedes idahonesis* was synonymized with *Ae. spencerii* by Wood [76] based on the lack of diagnostic characters for distinguish both species in larval and male stages. However, *Ae. idahoensis* continued recognized as a subspecies of *Ae. spencerii* for some authors [71, 77]. Finally, *Ae. idahoensis* was recognized as a valid species based on differences of adult’s scale patterns on abdominal ornamentation and geographic distribution [78]. In Mexico, *Ae. spencerii* was originally reported in Aguascalientes by Vargas [61] in a list of Culicinae species occurring in Mexico; and in a second species list of mosquito species, Díaz-Najera and Vargas [62], reported *Ae. idahoensis* as valid species in Aguascalientes, omitting the previous *Ae. spencerii* record. The presence in Mexico of *Ae. idahoensis* as *Ae. sepencerii* was repeated in subsequent mosquito species catalogs by Darsie [79] and Ortega-Morales et al. [80]. The specimens originally reported as *Ae. idahonensis* in Aguascalientes were found in the CAIM collection: four microscope slides (Catalog number: CAIMCilH/lam-03522–03525) containing one fourth instar larvae each in Canada Balsam as mounting medium, and each slide with curatorial data on right label: Number for all microscope slides 3163; Collection site: Aguascalientes City, Aguascalientes state; presumably collected on 21-Agoust-1941; by A. Martínez-Palacios: Specimens were photographed by H.H. and re-identified based on external characters by A.I.O.-M. as a mixed species of *Ae. nigromaculis* and *Ae. muelleri* [71]. CAIMCulH/lam-03522: *Ae. nigromaculis*; 03523: missing structures, possibly *Ae. muelleri*; 03524–03525: *Ae. muelleri*. Since the only record of *Ae. idahoensis* reported as *Ae. spencerii* in Mexico is those from Aguascalientes, and because the specimens from those records were re-identified as *Ae. muelleri* and *Ae. nigromaculis*, we recommend that *Ae. idahoensis* is removed from the list of Mexican mosquito species.

### Medically important mosquitoes of Aguascalientes

Some species reported in Aguascalientes are of medical and veterinary importance because they are vectors of pathogens causing diseases. Table 6 shows the most important public health species that occur in Aguascalientes. Host-seeking species caught by human landing methods and/or Shannon traps with humans were noted.

## Acknowledgments

We thank Salvador Morales-Avitia for his valuable collaboration during field surveillance studies. We thank Abdisalam Abdi, Yuexun Tian, and Juan Pablo Fimbres Macias for supporting samples.

## Author Contributions

**Conceptualization:** Aldo I. Ortega-Morales, Isis J. Morales-Avitia, Guadalupe Flores-Salas

**Data curation:** Aldo I. Ortega-Morales, Herón Huerta, Nadia A. Fernández-Santos, Isis J. Morales-Avitia, Guadalupe Flores-Salas

**Formal analyses:** Nadia A. Fernández-Santos, Lihua Wei, Miguel Medrano-Santillana, Sujata Balasubramanian, Gabriel Hamer

**Funding acquisition:** Nadia A. Fernández-Santos, Mario A. Rodríguez-Pérez, Jessica M. Flores-Salas, Jorge J. Rodríguez-Rojas

**Investigation**: Aldo I. Ortega-Morales, Isis J. Morales-Avitia, Guadalupe Flores-Salas, Jorge J. Rodríguez-Rojas

**Methodology:** Aldo I. Ortega-Morales, Jorge J. Rodríguez-Rojas

**Project administration**: Nadia A. Fernández-Santos, Mario A. Rodríguez-Pérez, Jessica M. Flores-Salas, Jorge J. Rodríguez-Rojas

**Resources**: Aldo I. Ortega-Morales, Nadia A. Fernández-Santos, Mario A. Rodríguez-Pérez, Jessica M. Flores-Salas, Jorge J. Rodríguez-Rojas, Miguel Medrano-Santillana, Sujata Balasubramanian, Sarah Hamer, Gabriel Hamer

**Software**: Miguel Medrano-Santillana, Sujata Balasubramanian, Sarah Hamer, Gabriel Hamer

**Supervision**: Aldo I. Ortega-Morales, Nadia A. Fernández-Santos, Mario A. Rodríguez-Pérez, Jorge J. Rodríguez-Rojas, Gabriel Hamer

**Validation**: Aldo I. Ortega-Morales, Nadia A. Fernández-Santos, Mario A. Rodríguez-Pérez, Jorge J. Rodríguez-Rojas, Gabriel Hamer

**Visualization:** Nadia A. Fernández-Santos, Jessica M. Flores-Salas, Aldo I. Ortega-Morales, Jorge J. Rodríguez-Rojas, Miguel Medrano-Santillana

**Writing–original draft preparation:** Aldo I. Ortega-Morales, Mario A. Rodríguez-Pérez, Sujata Balasubramanian

**Writing–review and editing:** Mario A. Rodríguez-Pérez, Jorge J. Rodríguez-Pérez, Nadia A. Fernández-Santos, Sarah Hamer, Gabriel Hamer

## Data Availability Statement

All mosquitoes (mounted on pins and / or microscope slides) collected in Aguascalientes state during this study were deposited in the Culicidae Collection (CC-UL), Parasitology Department, Universidad Autónoma Agraria Antonio Narro-Unidad laguna (Autonomous Agrarian University Antonio Narro laguna unit), Torreón, Coahuila, Mexico. Sequences generated from barcoding are available at NCBI (PZ832164–PZ832185, PZ832187–PZ832196). Data from metabarcoding of bloodmeals are at NCBI SRA (Bioproject PRJNA1491369 Biosamples SAMN61459855 to SAMN61459904).

